# A novel approach to *Caudoviricetes* taxonomy utilising whole proteome structure-structure comparison

**DOI:** 10.1101/2025.08.06.668922

**Authors:** Linda Smith, Thomas S.B. Schmidt, Igor Tolstoy, Timofey Skvortsov, Colin Hill, Andrey N. Shkoporov

## Abstract

Viral proteins often evolve so rapidly that sequence similarity is lost even among conserved functional genes, complicating phylogenetic analysis and classification. This challenge is acute for *Caudoviricetes*, a highly diverse class of dsDNA tailed bacteriophages recently redefined by the International Committee on Taxonomy of Viruses (ICTV). To address the limitations of sequence-based taxonomy, we present a structure-based classification framework using whole-proteome structural clustering. From 4,082 exemplar genomes, we predicted 445,098 protein structures with ESMFold and clustered them using Foldseek. Genomes were encoded as binary profiles of structural fold presence, and phylogenetic distances were inferred and normalized using Relative Evolutionary Divergence (RED). This yielded a revised structural taxonomy comprising 159 orders, 267 families, 502 subfamilies, and 1,189 genera. We also introduce *PhagePleats*, a Python tool for classifying novel phage genomes based on structural similarity. Our approach highlights the utility of protein structure for resolving distant viral relationships.

## Introduction

Viral taxonomy remains one of the most challenging frontiers in biology. Unlike cellular organisms, which share universally conserved genes, no single genetic marker is universally conserved across all viruses [1,2]. They are believed to have evolved independently at least six times [3]; a concept reflected in their classification into six distinct viral realms. These polyphyletic origins make it fundamentally impossible to construct a single, unified phylogeny—or “megataxonomy”—that encompasses all known viruses [1]. Viral genomes— especially those of dsDNA tailed bacteriophages and archaeal viruses belonging to *Caudoviricetes*—are mosaics composed of exchangeable modular gene cassettes acquired through horizontal gene transfer [4–6]. This genomic patchwork further erodes clear evolutionary signals, complicating phylogenetic inference even among closely related phages [7].

Even individual viral proteins—such as tail fibers—can be chimeric in origin, further confounding homology detection [8]. As a result, a large proportion of predicted viral ORFs remain unannotated, contributing to the vast “viral dark matter” [9,10]. The combination of rampant mosaicism and rapid evolution highlights the difficulty of applying sequence-based approaches for resolving deep viral relationships. This is exemplified by *Caudoviricetes* viruses, the most abundant and diverse biological entities on Earth [11-13] with major ecological importance across different ecosystems [14-16]. As metagenomic efforts continue to collect an ever-growing catalogue of viral genomes, the challenge of assigning higher-order taxonomic labels remains largely unresolved. In 2022, tailed bacteriophage taxonomy was overhauled with the establishment of the class *Caudoviricetes* and the dissolution of the morphology-based order *Caudovirales* and its families Myoviridae, Podoviridae, and Siphoviridae [17]. This revision enabled the creation of new orders based on genomic similarity, beginning with *Crassvirales* [18,19], followed by others including *Autographivirales* and *Pantevenvirales*. Despite this progress, the current ICTV taxonomy remains incomplete— only 24.5% (1,429 of 5,822) of *Caudoviricetes* species in the ICTV VMR40 (Virus Metadata Resource, release 40) have been assigned to an order and 58.2% (3,393 of 5,822) a family.

We posit that structure-based approaches to viral classification can mitigate issues faced by sequence-based viral phylogenies. Protein secondary and tertiary structures, constrained by function, tend to evolve more slowly than primary structure i.e. amino acid sequences [20], making them not only stable molecular markers of deep evolutionary or functional relationships, but phylogenetic relics encoding a historical record of divergence. Tools such as AlphaFold and ESMFold now enable high-throughput prediction of protein structures with near-experimental accuracy [21-23]. The ESM-2 model, for example, has been used to generate over 600 million predicted structures, constituting the first large-scale structural metagenomic protein database. Concurrently, tools like Foldseek [24] allow for ultrafast structural comparisons by encoding protein geometry into a 3Di alphabet, making it computationally feasible to search for conserved folds across millions of proteins using sequence comparison algorithms.

The growing availability of tools for large-scale structural prediction and comparison marks a timely inflection point for virology and phylogenetics. While the concept of structure-based viral taxonomy is not new [25,26], recent advances now make it feasible to apply these approaches systematically across the whole structurome of entire viral clades. In cases where sequence-based methods reach their limits, structural conservation offers a robust alternative, suggesting that structure-guided analyses may increasingly complement — and in some contexts, surpass — sequence-based strategies as a standard approach to viral classification. While prior efforts have used structural comparisons of single hallmark proteins to infer viral phylogeny [27], our approach extends this principle to the entire structural proteome. This offers a more holistic and evolutionarily relevant framework for taxonomic classification. To illustrate this, consider a hypothetical scenario: if a viral genome were entirely recoded to use different codons or alternative amino acid sequences, yet all resulting proteins retained their tertiary and quaternary structures, the virus would likely retain its original morphology, infectivity, and ecological role. From a functional and ecological perspective, it would still belong to the same lineage — despite being unrecognizable to sequence-based approaches.

Here we present a unified, whole-structurome–based taxonomic approach to resolve higher-order classifications within *Caudoviricetes*. We re-organize *Caudoviricetes* diversity along genome-wide structural similarity partitions that align remarkably well with established sequence-based ICTV taxonomic groups, while adding further resolution and establishing novel groups relative to existing classifications. Normalizing derived structural “phylogenies” by the *relative evolutionary divergence* [RED] among viral genomes, we delineate robust and reproducible clusters at different taxonomic depths. Finally, we introduce PhagePleats, a machine learning-based tool that accurately classifies novel genomes against the resulting taxonomies based on folded viral proteins.

## Results

### Deriving a Structure-Informed Strategy

A total of 4,082 genomes were selected from the ICTV VMR40 dataset to ensure representation from all recognized orders within *Caudoviricetes*. Currently, the class comprises 11 orders, of which 7 contain fewer than 10 members (Supplementary Figure 1, Panel A). Among the selected genomes, 3,291 out of 4,082 species lacked an order-level assignment; while across the entire *Caudoviricetes* dataset, 4,392 species remain unassigned at the order level. A similar pattern is observed at the family level: 2,257 genomes in the training set and 2,427 genomes overall lack a family designation (Supplementary Figure 1, Panel B). These figures highlight the limited resolution of current higher-order taxonomic groupings and underscore the need for a system that enables consistent and scalable classification at the order and family levels.

Given that tailed dsDNA bacteriophages and archaeal viruses (*Caudoviricetes*) share a conserved set of core structural proteins involved in capsid morphogenesis and DNA packaging, including the HK97-like major capsid protein (MCP), the portal protein superfamily (e.g., SPP1 GP6-like), and the terminase large subunit (TerL) superfamily, structural clustering was performed with the expectation that a sufficiently sensitive approach would group each of these protein superfamilies into distinct clusters, ideally with a representative present in the majority of genomes. To balance sensitivity and specificity in Foldseek clustering, parameter sets were benchmarked using three conserved viral proteins - major capsid protein (MCP), terminase large subunit (TerL), and portal - as internal controls. As shown in Supplementary Figure 2, increased sensitivity yielded larger clusters but reduced functional specificity (measured by Shannon entropy of predicted Pfam annotations), while higher specificity often led to fragmented clusters. Optimal runs were defined as those with a median cluster size ≥ 3,000 across MCP, TerL, and portal clusters, and a functional specificity ≥ 0.7. Structural coherence was further validated using PyMOL by computing the median and standard deviation of RMSD values across the five largest clusters (Supplementary Figure 3). The best-performing parameter set—coverage 40%, TM-score 30%, e-value 1E-2— yielded the largest median cluster size (3,367 across the top five largest clusters) and the lowest median RMSD (4.03), and was therefore selected for downstream analysis.

### Structural Conservation and Relatedness within *Caudoviricetes*

The most conserved structural protein across *Caudoviricetes* is the major capsid protein (MCP), with an average of 89% of genomes per order containing a homolog within the MCP cluster (1. *Phage major capsid protein E* | *YP_010082820*.*1*; Supplementary Figure 4). An exception is *Crassvirales*, where only 1% of genomes match this cluster; instead, their MCPs align with a distinct structural cluster—*Major virion structural protein Mu-1/Mu-1C (M2)*— highlighted in Supplementary Figure 4A (2). The terminase large subunit was the next most conserved, with approximately 75% of genomes per order containing a homolog in cluster 3 (*Terminase RNaseH-like domain* | *YP_009620569*.*1*). The tail tube protein, represented by cluster 5 (*Phage tail tube protein* | *YP_008058468*.*1*), also showed broad conservation, with most orders containing homologs. However, in *Autographivirales, Crassvirales*, and *Grandevirales*, tail tube proteins were more closely related to cluster 7 (*Tail tubular protein* | *YP_009098374*.*1*), indicating structural divergence in these groups.

To identify the most conserved functional domains across *Caudoviricetes* orders, the Pfam with the highest median representation was selected within each functional category. The most widespread domain overall was a protein of unknown function, DUF1128 (PF06569), present in an average of 77% of genomes per order and described as a family of small, uncharacterized bacterial proteins. Given its high conservation across *Caudoviricetes* orders suggests an important but currently uncharacterized role in the viral lifecycle, possibly related to genome regulation or host interaction.

Among annotated functional categories, the most conserved domains included a regulatory protein (YozE SAM-like fold, PF06855), potentially involved in DNA binding or signaling; a replication-associated component (TFIIH subunit Tfb5, PF06331), part of the transcription initiation complex; and a packaging-related domain (terminase small subunit, PF07141), essential for genome encapsidation. Each of these were present in over 40% of genomes on average at the order level. Moderately conserved functions (∼35%) included metabolic enzymes (6-carboxyhexanoate–CoA ligase, PF03744), likely involved in fatty acid or CoA metabolism, and host interaction proteins (G protein spike, PF02306), mediating viral entry. Less frequent but broadly distributed domains included those involved in structure (virus coat protein, PF00721), lysis (hemolysin XhlA, PF10779), defense (Clostridium-like enterotoxin, PF03505), and mobile genetic elements (HNH endonuclease, PF13392. These patterns underscore the conserved nature of replication and packaging machinery across *Caudoviricetes*, while also highlighting the potential biological significance of accessory genes that remain poorly characterized. Their consistent presence across diverse taxa suggests that key aspects of viral function and evolution are yet to be fully elucidated, warranting further investigation into these enigmatic components of the viral genome.

To quantify relatedness within and between taxonomic ranks, pairwise percentages of shared structural proteins were calculated for each clade (intra-rank) and between clades (inter-rank). Intra-clade similarity followed the expected trend, increasing toward lower taxonomic ranks with genus-level clades sharing an average of 55.7% ± 3.5%, followed by subfamily (44.9% ± 8.4%), family (37.5% ± 10.6%), and order (32.8% ± 12.1%). In contrast, protein sharing between clades was uniformly low across all ranks, with mean inter-clade similarity ranging from 4.8–5.2%, and correspondingly low variability (e.g., standard deviation = 0.5% at Genus, 1.5% at Order), indicating well-defined boundaries between taxonomic groups (Supplementary Figure 5).

### Structure-based taxonomy aligns with ICTV classifications and highlights gaps in higher taxonomic ranks

To assess how structural similarities among *Caudoviricetes* reflect existing taxonomy, a neighbour-joining phylogeny was constructed from a presence–absence matrix of structural protein clusters across 4,082 genomes, plus one outgroup. Herpes simplex virus 1 (HSV-1), a distantly related member of *Herpesviricetes* and a suitable outgroup to *Caudoviricetes* - as both classes belong to the realm *Duplodnaviria* - was selected as the outgroup for rooting the phylogram. Each genome was encoded as a binary vector denoting the presence or absence of each cluster, and pairwise Dice distances were used to generate the tree using DecentTree. The resulting structure-based phylogram showed majority agreement with ICTV taxonomy at both the family and order levels (Figure 2), supporting its use as a framework for resolving relationships within classified groups and guiding the classification of unassigned taxa.

**Figure 1.**
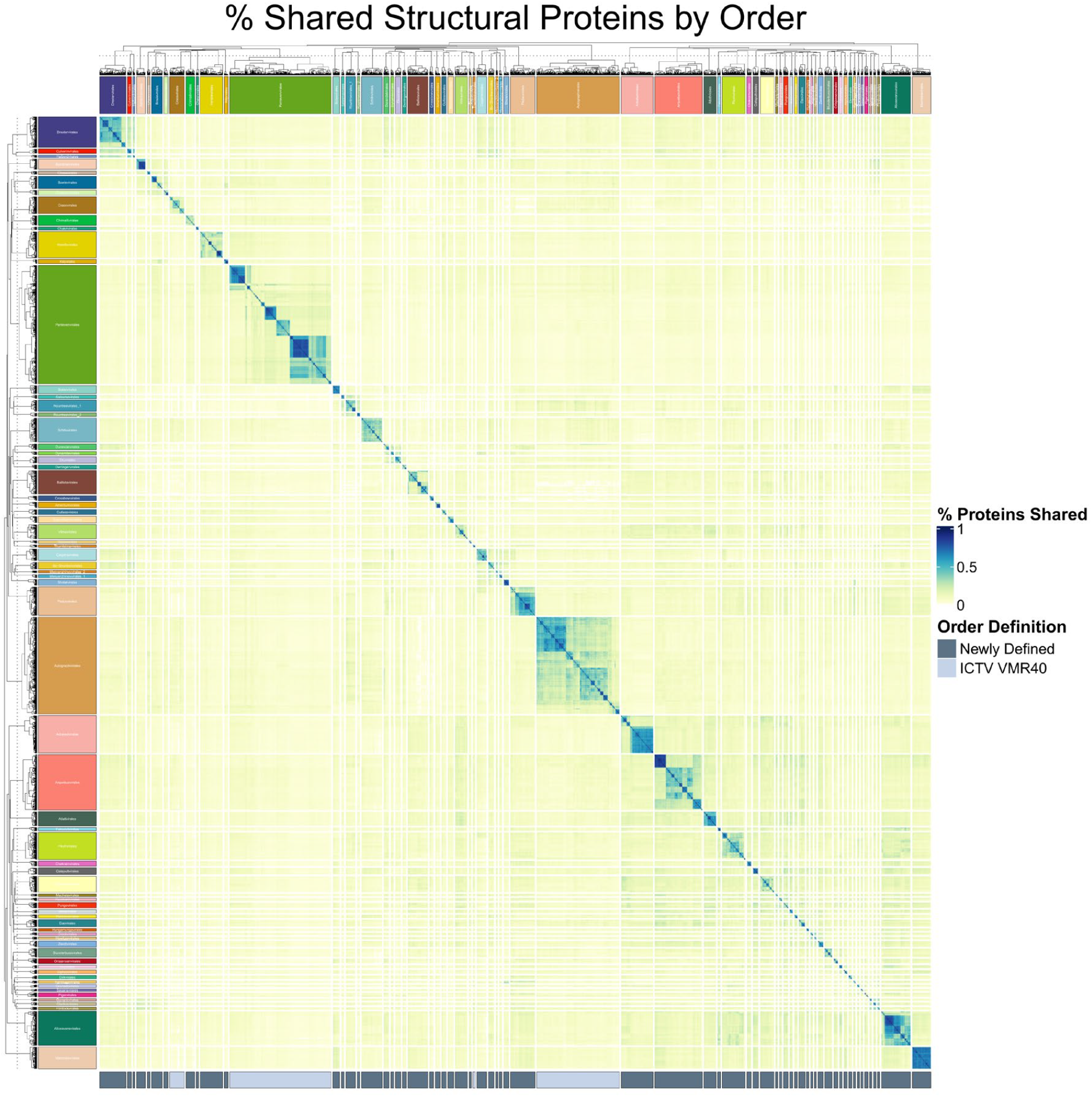
Structural protein similarity between viral genomes clustered by Order. The bottom panel compares order-level annotations derived from the newly defined RED-based taxonomy (see below) versus existing family assignments in ICTV VMR40. Orders with ≥ 10 members are included in the plot.

**Figure 2.**
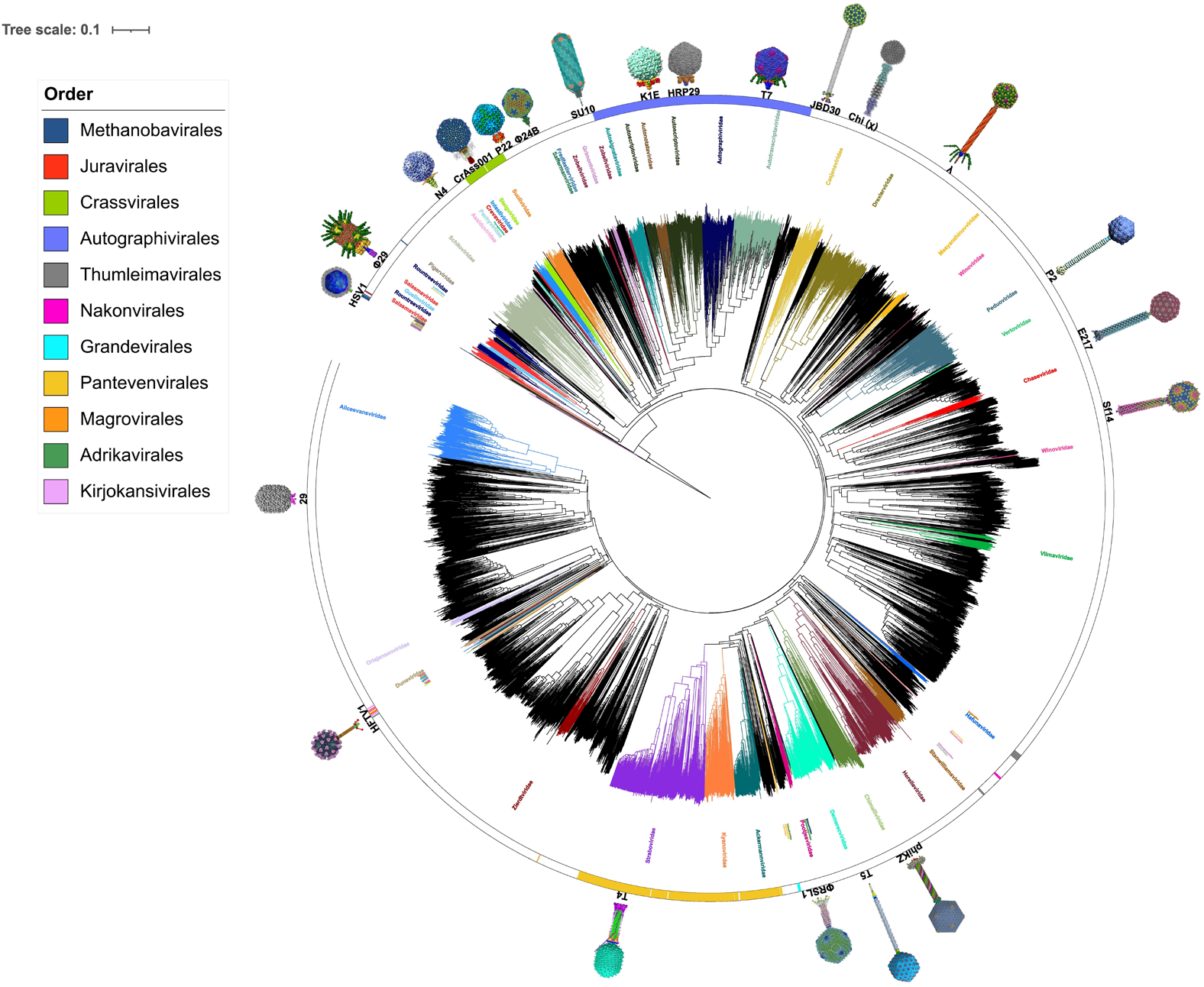
Structure-based cladogram of *Caudoviricetes* annotated with ICTV VMR40 taxonomy and cryo-EM-resolved representatives. A phylogram constructed from pairwise Dice distances between genomes based on the presence–absence of structural protein clusters; branch lengths reflecting global structural dissimilarity is shown. Each leaf is coloured by its ICTV-assigned family, with outer ring annotations indicating order-level classifications. A phylogram with the same topology, but with branch lengths normalized for uniformity, is shown in Supplementary Figure 6.

**Figure 3.**
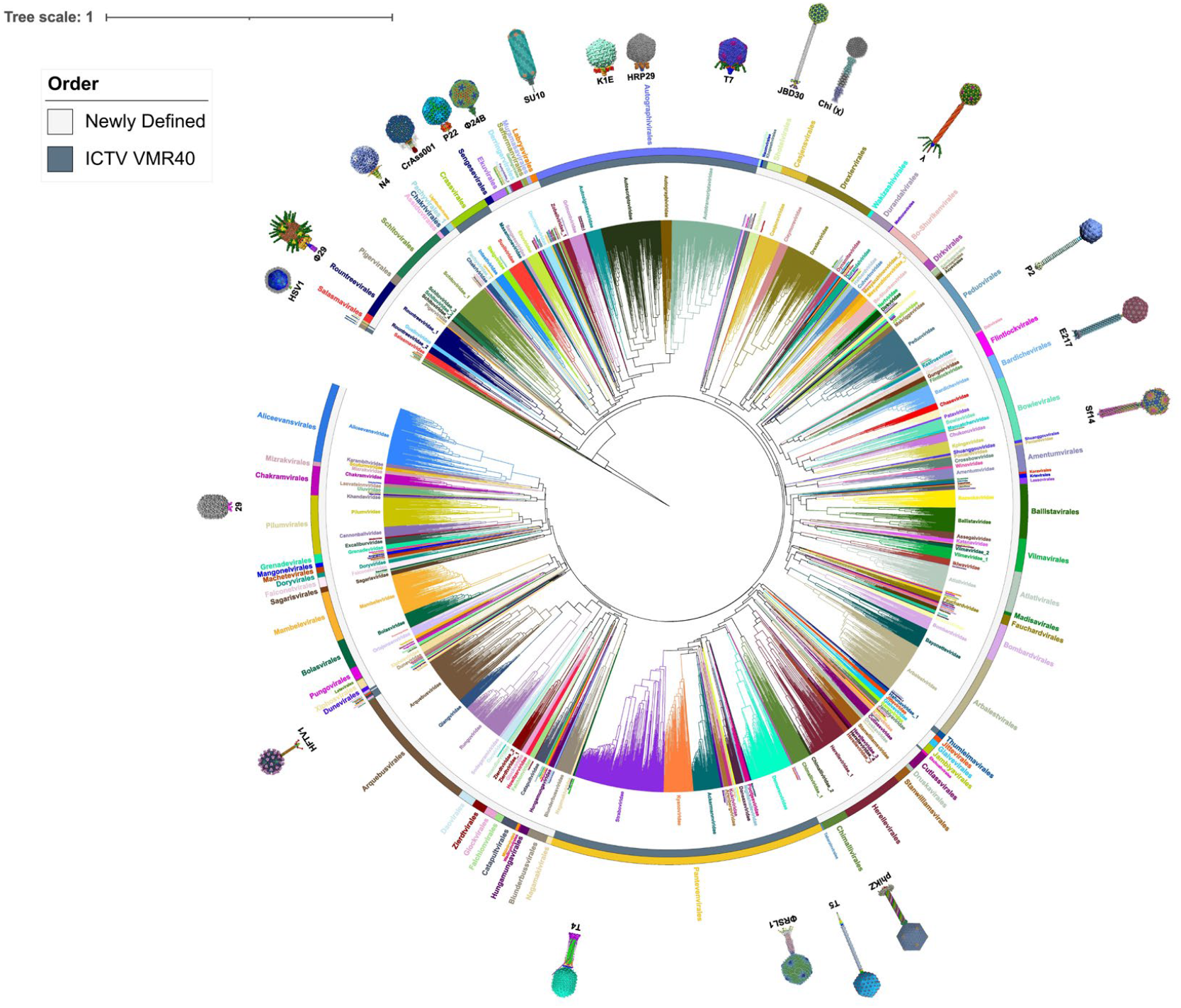
RED-scaled phylogenetic tree of *Caudoviricetes* displaying ICTV-recognized and newly defined orders and families. Rooted on the outgroup *Herpes simplex virus 1* (HSV-1), the circular cladogram depicts structural phylogenetic relationships among *Caudoviricetes*, scaled by Relative Evolutionary Divergence (RED). Branches are coloured by family-level classification, while the outer ring denotes order-level assignments. Orders recognized by ICTV VMR40 are indicated dark grey on the first outer ring.

The extent to which structural clustering reflects ICTV taxonomy was quantified by assessing monophyly across taxonomic ranks, using the last common ancestor (LCA) of each group in the structural phylogram. For each ICTV-assigned taxon containing two or more genomes, a precision score was computed, defined as the proportion of genomes within the last common ancestor (LCA) that were assigned to the same taxonomic rank. A minimum of two genomes were required to assess monophyly, as a single genome cannot form a clade or reveal whether related genomes cluster together. Among the 80 ICTV-recognized families in the dataset, 55 met this criterion and were included in the analysis.

Among the 55 assessed families, 30 (54.5%) were perfectly monophyletic (precision = 1.0), and 38 (69.1%) achieved precision scores above 0.85, indicating strong concordance between structural clustering and ICTV family assignments. Perfect monophyly was observed in both large families (e.g., *Demerecviridae, Drexlerviridae, Herelleviridae, Peduoviridae*) and smaller ones (e.g., *Fredfastierviridae, Graaviviridae*). Near-perfect groupings included *Ackermannviridae* (precision = 0.984) and *Straboviridae* (0.981), with minor deviations resulting from the inclusion of previously unassigned genomes.

Inconsistencies were apparent for several families. *Rountreeviridae, Salasmaviridae*, and *Guelinviridae* were fragmented across neighbouring clades, with *Guelinviridae* embedded between two branches of *Salasmaviridae*, resulting in low precision scores (0.521, 0.352, and 0.172, respectively). Further conflicts emerged within the order *Autographivirales*, where *Autotranscriptaviridae* was nested within *Autographiviridae*, indicating a single monophyletic lineage of which *Autographiviridae* is the ancestral clade. Similarly, *Autonotataviridae* (precision = 0.275) appeared as a nested subcluster within *Autoscriptoviridae*, the dominant family in that region of the tree.

At the order level, several taxa also exhibited strong structural concordance. *Nakonvirales* and *Grandevirales* were fully monophyletic (precision = 1.0), while *Crassvirales* (0.973), *Pantevenvirales* (0.949), and *Autographivirales* (0.918) showed high precision scores, indicating substantial agreement with structure-based clustering. Minor deviations from perfect precision in these groups reflect the incorporation of previously unclassified genomes into the respective clades. In contrast, orders of archaeal viruses such as *Methanobavirales, Thumleimavirales, Kirjokansivirales, Magrovirales*, and *Juravirales* were highly fragmented, with precision scores ranging from 0.001 to 0.031.

Subfamily- and genus-level classifications showed the highest consistency with structural clustering. Of the 121 subfamilies present, 85 (70.2%) were perfectly monophyletic, and among 584 genera, 503 (86.1%) showed complete monophyly (precision = 1.0) (Supplementary Table 1, Supplementary Figure 7).

### Taxonomic Expansion Through Relative Evolutionary Divergence (RED) Scores and the definition of new candidate taxa

To resolve polyphyletic ICTV taxa, address ambiguous or incomplete classifications, and enable systematic taxonomic assignment across all genomes in the structural phylogeny, a topology-aware, rank-normalized framework was applied using Relative Evolutionary Divergence (RED) values. Relative Evolutionary Divergence (RED) is a phylogenomic metric that quantifies the relative position of each node along a rooted tree, with values scaled from 0 at the root (representing the most recent common ancestor) to 1 at the tips (extant taxa), therefore assigning a scaled position to each node based on its evolutionary distance from the root to the tips.

Internal nodes are assigned RED scores through linear interpolation based on branch lengths, using the formula: p + (d / u) × (1 – p), where *p* is the RED of the parent node, *d* is the branch length between node *n* and its parent, and *u* is the average branch length from the parent to all descendant tips of *n*. This approach is conceptually aligned with the method used by the Genome Taxonomy Database (GTDB) to delineate bacterial taxa [28]. Unlike approaches that rely on absolute branch length thresholds, RED enables the use of relative, rank-normalized cut-offs across taxonomic levels, allowing for a more scalable and reproducible classification framework.

To compute RED scores, the structural phylogeny - rooted on *Herpes simplex virus 1* (HSV- 1) - was processed using PhyloRank, which assigned RED values to all internal nodes and generated a RED-normalized tree suitable for downstream taxonomic standardization. Optimal RED thresholds for each taxonomic rank were determined by iteratively scanning RED cut-offs along the phylogenetic tree, from the root (0) to the tips (1), and assessing the agreement between RED-defined clades and ICTV taxonomy using the Adjusted Mutual Information (AMI) score - a chance-corrected metric of clustering similarity. AMI values for each taxa were plotted against RED thresholds (Supplementary Figure 8), and local maxima in the curve were interpreted as optimal divergence points for each rank. This approach yielded RED intervals with maximal alignment to ICTV taxonomic levels: Order (0.46–0.54), Family (0.54–0.63), Subfamily (0.63–0.77), Genus (0.77–0.92), and Species (0.92–1.00). These thresholds were then applied to resolve conflicts and assign taxonomic ranks throughout the tree, including those previously lacking classification. A visual representation of these RED-informed boundaries is provided in Supplementary Figure 9.

To evaluate the agreement between RED-defined clades and ICTV taxonomy, the proportion of taxa recovered as monophyletic at the optimal RED threshold for each rank was quantified, as determined by peak Adjusted Mutual Information (AMI) scores. A taxon was considered monophyletic if all genomes assigned to it clustered within a single RED-defined clade. At the order level (RED = 0.46), 55% of ICTV-defined orders were monophyletic (6/11). Family- and subfamily-level thresholds (RED = 0.54 and 0.63, respectively) yielded similar recovery rates, with 65 out of 79 families (82.3%) and 102 out of 124 subfamilies (82.3%) forming single clades. Genus-level recovery was highest, with 1,302 out of 1,367 genera (95.2%) recovered at RED = 0.77. These results highlight strong topological coherence between RED-informed clades and existing ICTV taxonomy, particularly at finer taxonomic ranks.

In cases where ICTV-defined taxa were identified as polyphyletic, RED-informed clade boundaries were used to guide reclassification. The largest monophyletic clade was retained under the original ICTV designation, while smaller, topologically inconsistent subclades were assigned novel placeholder names to maintain phylogenetic coherence. For example, the family *Herelleviridae* was partitioned into four distinct clades based on the RED-derived family-level threshold, resulting in designations such as “Herelleviridae_1”, “Herelleviridae_2”, and so on. Novel orders were named using the prefix of the majority family within each clade, whereas novel families were assigned names derived from classical weaponry, and novel subfamilies were named after natural satellites in the solar system.

At the order level, 146 novel orders were created, with an average of 20.7 genomes per order (median = 8, range = 1–244), including 3 singleton clades. At the family level, 200 new families were defined, with a mean size of 10.6 genomes (median = 4, range = 1–118), including 23 singletons. Subfamily-level definitions showed even greater granularity, with 356 novel subfamilies averaging 5.6 genomes each (median = 3, range = 1–95), including 96 singletons. This framework enabled systematic taxonomic expansion across the phylogeny, culminating in the proposal of a total 159 orders, 269 families, 503 subfamilies, and 1,189 genera within the dataset.

### Machine Learning-Based Taxonomic Classification

To extend this analysis into a predictive framework for assigning taxonomy to novel phage genomes, binary Random Forest classifiers were trained across the four hierarchical taxonomic ranks: order, family, subfamily, and genus. Each genome was encoded as a binary vector indicating the presence or absence of ∼20,000 non-redundant structural protein clusters, providing a representation of its structural proteome. Classifier training was supervised using labels derived from revised and extended taxonomic assignments based on Relative Evolutionary Divergence (RED) cut-offs.

The most predictive features for distinguishing taxonomic ranks were minor tail protein (YP_010055968.1), followed by two DNA polymerases (YP_009855449.1, YP_010062267.1), a DNA primase/helicase (YP_009824164.1), and a portal protein (YP_009802963.1). Of the top 30 most important features, 22 were associated with structural or replicative functions, consistent with these proteins playing central roles in *Caudoviricetes* biology and exhibiting lineage-specific diversity that enables taxonomic discrimination. Collectively, these top 30 features (Figure 5) accounted for approximately 15% of the total model’s feature importance, suggesting that a small subset of proteins disproportionately contribute to accurate taxonomic classification. These features reflect conserved functional and phylogenetic signatures, indicating that the models rely on biologically meaningful signals to distinguish viral taxa. Classifier performance across taxonomic ranks was evaluated as a function of training set size, with all taxa downsampled based on the distribution of member counts per taxon (Supplementary Figure 8) to ensure balanced class representation, while the test set was left unstratified to reflect the natural class distribution. At higher ranks (order and family), performance was consistently strong across all metrics - even with as few as 5–10 genomes per taxon - highlighting the robustness of structural features for coarse-level classification. AUROC, cross-validation, precision, and F1 scores all exceeded 0.95 at relatively low sampling depths (Supplementary Figure 11-14), suggesting that these broader clades are well-captured by the underlying structural signal. Notably, subfamily-level performance remained stable across training sizes (Supplementary Figure 13), suggesting that subfamily ranks may represent an optimal balance between taxonomic resolution and predictive robustness. At the genus level, all test metrics surpassed the 95% threshold at ≥10 training leaves but saw a gradual decline in precision (positive class) and F1-score (positive class) at training sizes ≥15, due to reduced representation of the positive class in the test set (Supplementary Figure 14).

**Figure 4.**
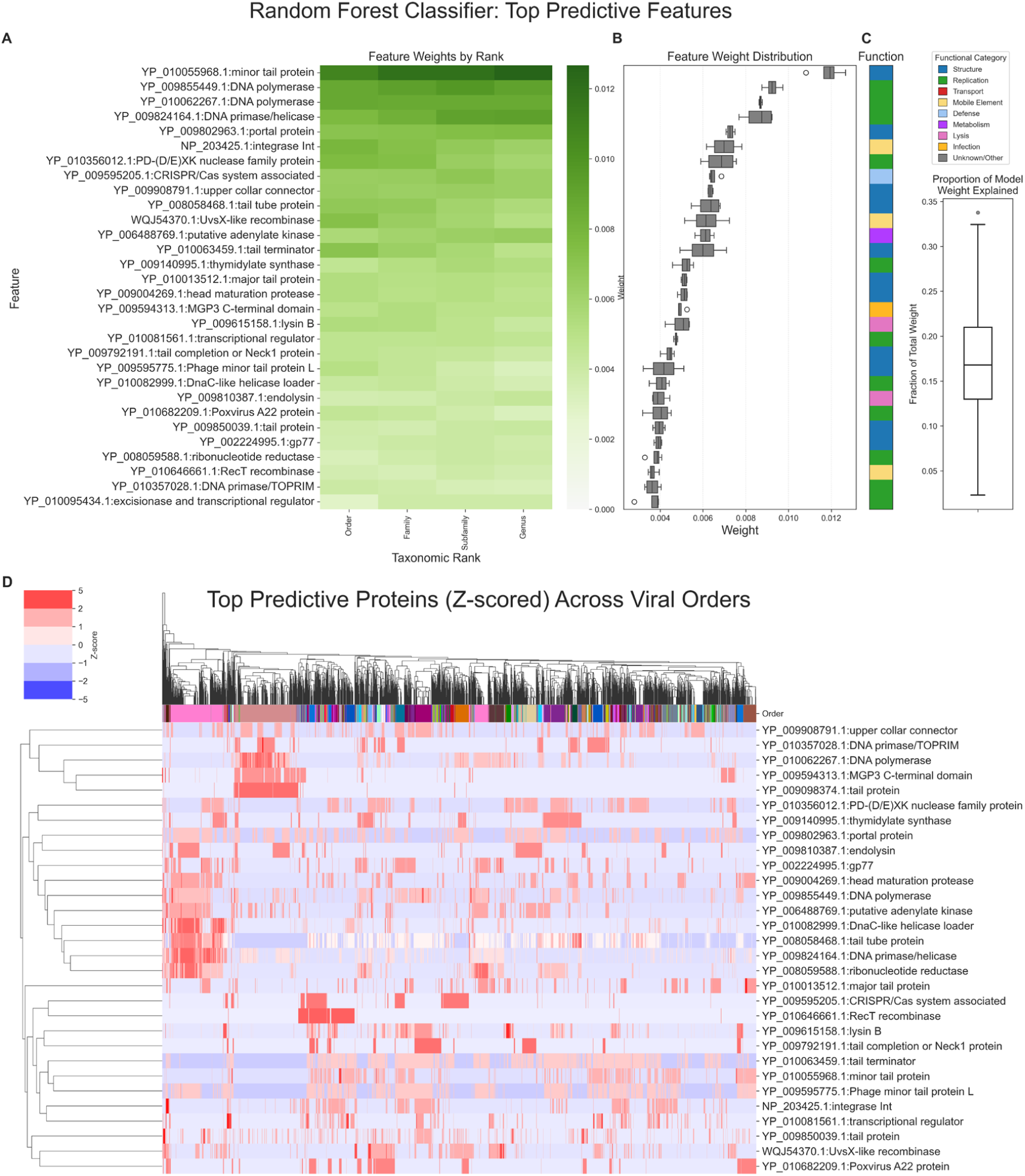
Top Predictive Protein Features Used by the Random Forest Classifier Across Viral Taxonomic Ranks. **(A)** Heatmap showing the feature weights (importance scores) assigned by the Random Forest classifier to individual proteins, across multiple taxonomic ranks (Order, Family, Subfamily, Genus, Species). Highly weighted features include structural proteins (e.g., tail proteins, portal proteins) and replicative enzymes (e.g., DNA polymerases, integrases), which are informative for classification across multiple ranks. **(B)** Boxplot of the distribution of feature weights across all models, highlighting the top predictive proteins ranked by median weight. **(C)** Functional annotations of each protein, colour-coded by category (e.g., Structure, Replication, Transcription, etc.). The vertical barplot on the right shows the proportion of total model weight explained by these top features, indicating a skew toward structural and replication-associated functions. **(D)** Hierarchically clustered heatmap showing the z-scored presence-absence or abundance profiles of the top predictive proteins across viral orders. Rows are proteins, and columns are viral genomes coloured by their predicted order. This view illustrates how different orders exhibit distinct protein usage patterns, reinforcing the discriminatory power of these features. The clustering highlights both order-specific markers and broader patterns of conservation or variability.

The final production classifier was trained on all available taxa without downsampling, maximizing its exposure to the full diversity of known *Caudoviricetes*. To evaluate performance on unseen data, the production models were tested on an independent set of 1,746 ICTV-classified *Caudoviricetes* genomes (131,003 proteins) that were excluded from training. Of the 603 genomes with order-level labels matching those in the training set, 602 (99.8%) were correctly classified, suggesting that the single misclassified genome may have been incorrectly labeled in the reference dataset rather than mispredicted by the model. At the family level, 785 out of 854 genomes (91.9%) were correctly predicted; 941 of 1,019 subfamily labels (92.4%) were accurate, and 998 out of 1,141 genus-level predictions (87.4%) matched their ground truth assignments (Supplementary Figure 15).

### PhagePleats: A Tool for Taxonomic Classification of Phages

This work culminated in the development of PhagePleats (github.com/linda5mith/PhagePleats), a Python tool for taxonomically classifying phage genomes based on their pre-folded proteomic structures, incorporating the methodology defined in this study. The tool uses a reference database of ∼20,000 structural protein clusters derived from the clustering output that underpins the structural phylogeny. For each input genome, its pre-folded proteins (in PDB format) are searched against this reference using Foldseek. Based on the resulting matches, a binary presence–absence vector is generated to denote which structural clusters are represented in the input genome. These vectors are assembled into a matrix and fed into the suite of binary Random Forest classifiers, where the highest probability taxa for each rank is returned.

To account for the immense unexplored diversity of the virosphere - and the likelihood of encountering highly divergent or novel tailed viruses - a novelty-aware post-prediction quality control layer was integrated into the pipeline. This consists of a nearest-neighbour similarity search for each genome using the FAISS (Facebook AI Similarity Search) library. For each input genome, the most similar genome in the training set is identified based on Jaccard similarity, which quantifies the percentage of shared structural protein clusters. These nearest-neighbour comparisons serve two key purposes. First, they provide a secondary validation layer for classifier predictions—particularly when model confidence is low. Second, they enable biologically interpretable validation by confirming that each input genome shares structural similarity with at least one training genome. For instance, if an input genome shares 0% of structural proteins with any genome in the training set, it can be confidently flagged as novel.

In addition, intra-clade statistics were computed for each taxonomic rank in the training set, including the mean and standard deviation of percentage shared proteins within each clade. These distributions were then used to calculate z-scores for each input genome’s similarity and distance to its predicted clade. A novelty flag was assigned based on the extremity of these z-scores compared to clade-level distributions—using thresholds of ±1, ±2, and ±3 standard deviations. Genomes were flagged as “Likely member”, “Potential new rank”, or “Novel taxa” if intra-clade statistics were unavailable. This approach allows PhagePleats to not only assign taxonomy, but also to quantify how divergent a query genome is from known lineages.

## Conclusion

We present PhagePleats, a Python-based framework for scalable, objective, and reproducible taxonomic classification of *Caudoviricetes* viruses based on structural proteome content. By integrating whole-structurome similarity, RED-based phylogenetic normalization, and machine learning, our approach achieves high congruency with existing ICTV taxonomy. Additionally, it expands the current classification system by assigning higher-level taxonomic ranks and delineating novel clades and lineages across the dataset. Importantly, this framework lays the foundation for a universal, structure-informed viral classification system capable of extending across the virosphere. As genomic databases continue to expand and the efforts of Viral Study Groups refine and formalize viral groupings, the integration of newly characterized genomes will further strengthen model performance and taxonomic resolution. With the ongoing growth of curated viral data, this approach holds the potential to assign taxonomy to all known and future viral genomes, bridging current classification gaps and supporting a unified, data-driven taxonomy for viral diversity.

## Methods

### Folding and Selection of Sample Genomes

A total of 4,082 representative *Caudoviricetes* genomes were selected from ICTV VMR40, ensuring broad taxonomic coverage across all orders (see 4083_ICTV_metadata.csv). One additional genome, *Herpes simplex virus 1* (HSV-1; NC_001806.2), was included as the outgroup for downstream tree rooting, bringing the total to 4,083 genomes. All 445,098 phage proteins (plus 53 HSV-1 proteins) were folded using the esmfold_v1() model from ESMFold (https://github.com/facebookresearch/esm). The environment was configured using the provided environment.yml file, with modifications for compatibility with an A100 GPU. Specifically, PyTorch was upgraded to torch==1.9.0+cu111, and the folding process was executed using fold.py with parameters --chunk-size 128.

### Structural Clustering, Protein Annotation and Quality Assessment

The 445,098 predicted protein structures were clustered using Foldseek’s easy-cluster module across a range of parameter combinations (execution scripted in run_foldseek_batches.sh). The resulting result.tsv files were imported into Notebook 1 (https://github.com/linda5mith/Structural_Taxonomy_Caudoviricetes/) for benchmarking. To identify optimal clustering parameters, multiple performance metrics were evaluated for each parameter set. These included: (i) median cluster size across known marker proteins - major capsid protein (MCP), terminase large subunit (TerL), and portal protein - required to be ≥ 3,000; and (ii) functional specificity ≥ 0.7, measured as the inverse Shannon entropy of Pfam domain annotations within each cluster. Foldseek easy-cluster runs that met these criteria were further evaluated for structural coherence. For each parameter set, the five largest resulting clusters were examined by computing the root-mean-square deviation (RMSD) of 3D structural superpositions between the cluster representative and its members using PyMOL (scripts/pymol_cluster_RMSD_top_N.py). Metrics calculated included: median RMSD per cluster, a summary statistic (median of the five cluster-level medians), and RMSD spread (standard deviation across member RMSDs). This benchmarking strategy ensured that clusters reflected both functional and structural homology. Based on these results, the optimal clustering parameters were selected as: coverage = 40%, TM-score = 0.30, E-value = 1E-2, cluster-reassign = 1. These settings yielded the largest median cluster size (3,367 members) and lowest median RMSD (4.03 Å) (Supplementary Figure 3).

Cluster members were annotated by linking RefSeq protein definitions, genome accessions, and ICTV-derived phage taxonomy to a manually curated protein_metadata.csv using in-house Python scripts (see *Notebook 1*). A total of 249 truncated TerL proteins were identified and excluded (terls_excluded_accns.csv). Truncated proteins were first flagged via visual inspection in PyMOL, where apparent half-length proteins were observed forming cluster representatives, and then formally filtered by excluding sequences shorter than half the median TerL length (273 amino acids, based on a median of 547aa across clusters). This step ensured removal of partial sequences that could bias clustering or functional annotation.

### pLDDT Threshold Justification and Fold Quality Evaluation

Although ESMFold provides global pLDDT scores as a proxy for structure confidence, no pLDDT-based filtering was applied. To justify this, we examined structural alignments from the three largest known clusters - terminase subunit (YP_009620569.1), MCP (YP_010082820.1), and tail tube (YP_008058468.1) - across a range of pLDDT values. Foldseek visualizations (--format-mode 3) showed that proteins with low global pLDDT scores (<70) still retained clear structural homology to cluster representatives, provided the TM-score threshold was ≥ 0.30. This confirmed that Foldseek alignments were robust even for modest-confidence folds. Additional exploration of Foldseek alignment and pLDDT relationships was examined in Notebook 2, and visual outputs were stored in data/pLDDT_vs_TM_score_alignment_html/^*^.html. Overall, pLDDT score skewed toward higher confidence, with most values falling between 70 and 90 (Supplementary Figure 13).

### Pfam Domain Annotation

To supplement existing RefSeq annotations, Pfam domain predictions were obtained using protein-vec (https://github.com/tymor22/protein-vec), which maps protein sequences to Pfam domain definitions via embedding-based similarity. The protein-vec environment was built using its provided environment.yml and adapted for CUDA 11.4 compatibility. This involved installing CUDA and PyTorch with: conda install cuda -c nvidia/label/cuda-11.4.0, conda install pytorch==1.12.1 torchvision==0.13.1 torchaudio==0.12.1 cudatoolkit=11.3 -c pytorch. Predicted Pfam accessions were linked to domain definitions from the Pfam-A seed database (https://www.ebi.ac.uk/interpro/download/Pfam/Pfam-A.seed.gz) using custom Python scripts (data/scripts/protein-vec/).

### Structural Presence–Absence Matrix and Protein Similarity (Sørensen– Dice Coefficient)

To quantify structural similarity between viral genomes, a binary presence–absence matrix of structural protein clusters was generated from the optimal Foldseek easy-cluster output (parameters: coverage = 40%, TM-score = 0.3, E-value = 1E-2, cluster-reassign=1). This clustering output was merged with protein metadata, allowing each protein to be annotated with its structural cluster, functional label, and source genome. A unique identifier was created by concatenating the cluster ID with its associated function. The merged dataset was converted into a binary matrix, with rows representing unique structural protein clusters (annotated by function) and columns corresponding to individual genomes. A value of 1 was assigned to a matrix entry if the genome contained a homologous protein belonging to the respective structural cluster, and 0 otherwise. Clusters present in fewer than two genomes were excluded to remove singletons from downstream analyses. To evaluate pairwise similarity between genomes, Sørensen–Dice similarity coefficients were computed for all genome pairs (Notebook 2). This metric quantifies the proportion of shared protein clusters between two genomes, normalized by the average number of clusters present in each. Specifically, for genomes A and B, Dice similarity was calculated as:

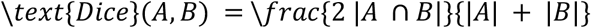

This resulted in a symmetric genome-by-genome matrix, where each value reflects the percentage of shared structural clusters. The Dice similarity matrix was converted to PHYLIP format and passed to DecentTree (decentree --in presence_absence.phylip --t NJ --out presence_absence.nwk) for phylogenetic reconstruction using the Neighbor-Joining (NJ) algorithm. The resulting tree was rooted on Herpes simplex virus 1 (HSV-1; NC_001806.2) and used in subsequent RED scaling and taxonomic classification analyses.

### RED-Based Phylogenetic Scaling and Taxonomic Boundary Inference

To assign relative evolutionary divergence (RED) values and define taxonomic boundaries in a phylogenetically informed manner, the presence–absence-based tree generated by DecentTree was rooted using the Herpes simplex virus 1 (HSV-1; NC_001806.2) outgroup. The rooted tree was passed to PhyloRank for RED score calculation and tree normalization using the following command: phylorank outliers --fixed_root -t phylorank_taxa_outgroups.txt --verbose_table 4083_rooted.nwk output_dir. The phylorank_taxa_outgroups.txt metadata file is generated in Notebook 3. The resulting RED-scaled tree and node metadata were saved as 4083_outgroup_rooted.scaled.tree and 4083_outgroup_rooted.scaled_rooted.node_rd.tsv, respectively.

These outputs provided a RED score for every internal node, which was then merged with genome-level metadata in Notebook 3. Taxonomic boundaries at the order, family, and subfamily levels were inferred by computing optimal RED thresholds that maximized Adjusted Mutual Information (AMI) scores when comparing RED-based clades to existing ICTV classifications. AMI, a measure of clustering agreement that adjusts for chance, was calculated across a range of potential RED cut-offs. The RED threshold corresponding to the maximum AMI for each rank was selected as the default boundary for defining monophyletic clades. These RED-informed boundaries were applied to the full *Caudoviricetes* tree to define consistent, rank-specific clades. Core clades for each taxonomic rank were established using Average Mutual Information (AMI) scores and RED thresholds (e.g., Order: 0.46–0.54), with a ±0.03 buffer to allow inclusion of borderline clades. For ranks such as Order and Family, these edge cases were visualized in iTOL and manually inspected to avoid excluding evolutionarily meaningful groups due to small deviations in RED (see Notebook 3).

These RED-informed boundaries were then applied to the full tree to define consistent, rank-specific clades across the *Caudoviricetes* dataset. Once core rank-level clades were established using Average Mutual Information (AMI) scores and RED-based thresholds (e.g., for Order, RED range = 0.46–0.54), a small buffer of ±0.03 was permitted to capture clades bordering the cut-off. In the case of order and family ranks, these minority borderline clades were visualized in iTOL and assessed manually to avoid excluding evolutionarily meaningful groups due to slight deviations in RED (see Notebook 3). To reduce redundancy, we applied a greedy algorithm that selects a minimal, non-overlapping set of Most Recent Common Ancestors (MRCAs). An MRCA here is defined as a monophyletic node in the tree with a RED score within the range for a given rank. The algorithm prioritizes lower RED scores and maximizes the number of unique descendant leaves while avoiding overlap. MRCA candidates were sorted by ascending RED and selected iteratively based on how many new, non-overlapping leaves they contributed. Clades slightly outside the target RED range but with sufficient exclusivity and size were flagged for manual review and included if justified. The resulting set of non-redundant, rank-informed MRCAs formed the basis for downstream classification.

### Random Forest Classifier Training

Binary “one-vs-rest” Random Forest classifiers were trained to predict taxonomic assignments at the Order, Family, Subfamily, and Genus levels using the RED-informed clades defined in Section 4 (see Notebook 4). Genomes were encoded as binary presence-absence vectors of clustered structural proteins. For each rank, taxa were modelled independently, with the goal of maximizing precision and recall across unbalanced classes. To evaluate model robustness under constrained data conditions, classifiers were trained across a range of balanced training sizes (e.g., 5, 10, 20, 50, 100 genomes per taxon). Only taxa with sufficient genome representation were included at each training size, and all included taxa were downsampled to ensure class balance. Taxa with fewer than five positive examples were excluded from cross-validation to maintain meaningful and stable fold splits. Classifier performance was evaluated across training sizes using multiple metrics, including cross-validation accuracy, AUROC, average precision (AUPRC), recall, precision and F1 score. Feature importance scores were recorded for each taxa rank and training size to identify consistently informative protein features.

Following benchmarking, final production classifiers were trained on the full dataset without downsampling or data splitting (train_production_classifiers() in Notebook 4). To assess their generalization, these models were evaluated on a held-out test set comprising genomes with ICTV-curated taxonomic labels that matched training-set clades. Any test genomes assigned to taxa not present in the training set (e.g., novel families) were excluded from evaluation to avoid penalizing the model for unclassifiable inputs. To validate classifier performance on unseen data, 1,746 ICTV-curated *Caudoviricetes* genomes not included in the training set were downloaded, and their proteins were folded using the same pipeline. Of these, 603 had *order-*level labels, 854 had *family-*level labels, 1,019 had *subfamily-*level labels, and 1,141 had *genus-*level labels (see Notebook 4). The production models generated in Notebook 4 are used in PhagePleats to predict an input genome’s taxonomy. All code and output data supporting this study are openly available in the GitHub repository: (https://github.com/linda5mith/Structural_Taxonomy_Caudoviricetes/).

## Acknowledgements

Linda Smith was funded by Research Ireland Centre for Research Training in Genomics Data Science [18/CRT/6214]. Andrey Shkoporov was funded by Wellcome Trust Research Career Development Fellowship [220646/Z/20/Z].This research was funded in whole, or in part, by the Wellcome Trust and was conducted with the financial support of Research Ireland under Grant Number 12/RC/2273-P2. For the purpose of open access, the authors have applied a CC BY public copyright license to any Author Accepted Manuscript version arising from this submission.

## Author contributions

Conceptualization, L.S., I.T. and A.N.S; Methodology, L.S., T.S.B.S., I.T., T.S. and A.N.S.; Investigation, L.S.; Writing – Original Draft, L.S.; Writing – Review & Editing, all authors; Funding Acquisition, L.S. and A.N.S.; Resources, I.T., T.S., C.H. and A.N.S.; Supervision, C.H. and A.N.S.

## Data availability statement

All data and code used in this study are publicly available at Zenodo (DOI: 10.5281/zenodo.15921490) and on GitHub at https://github.com/linda5mith/Structural_Taxonomy_Caudoviricetes.

