## Supplementary Figures for "A novel approach to *Caudoviricetes* taxonomy utilising whole proteome structure-structure comparison"

### Supplementary Material for: A Novel Approach to *Caudoviricetes* Taxonomy via Whole-Proteome Structure Comparison.

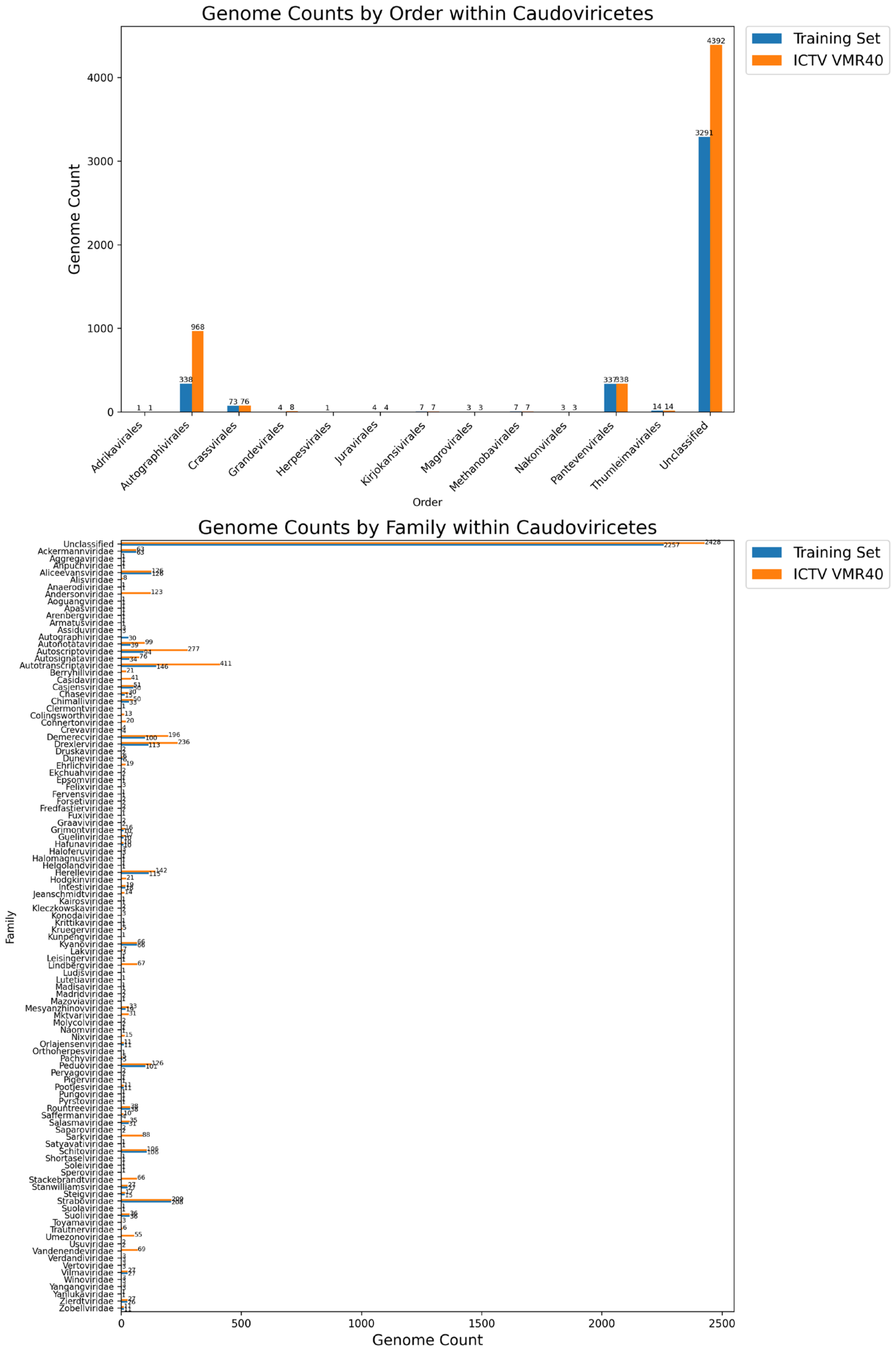


**Supplementary figure 1. Taxonomic representation of *Caudoviricetes* genomes in the training set versus the ICTV VMR40 dataset.**

**(A)** Bar plot showing genome counts by order for both the training set (blue) and the complete ICTV VMR40 dataset (orange). While orders such as *Autographivirales, Crassvirales and Pantevenvirales* are well-represented, the majority of genomes remain unclassified at the order level in ICTV VMR40. **(B)** Bar plot showing genome counts by family, highlighting the uneven representation of families within *Caudoviricetes*. Several families are highly enriched—such as *Autotranscriptaviridae*, *Demerecviridae*, *Drexlerviridae*, *Schitoviridae and Straboviridae*—each with over 100 members, while many others are sparsely represented or remain entirely unassigned. This highlights the absence of higher-level taxonomic definition within the current *Caudoviricetes* classification scheme (ICTV VMR40) and underscores the need for an expanded classification framework to capture the diversity of tailed dsDNA viruses.

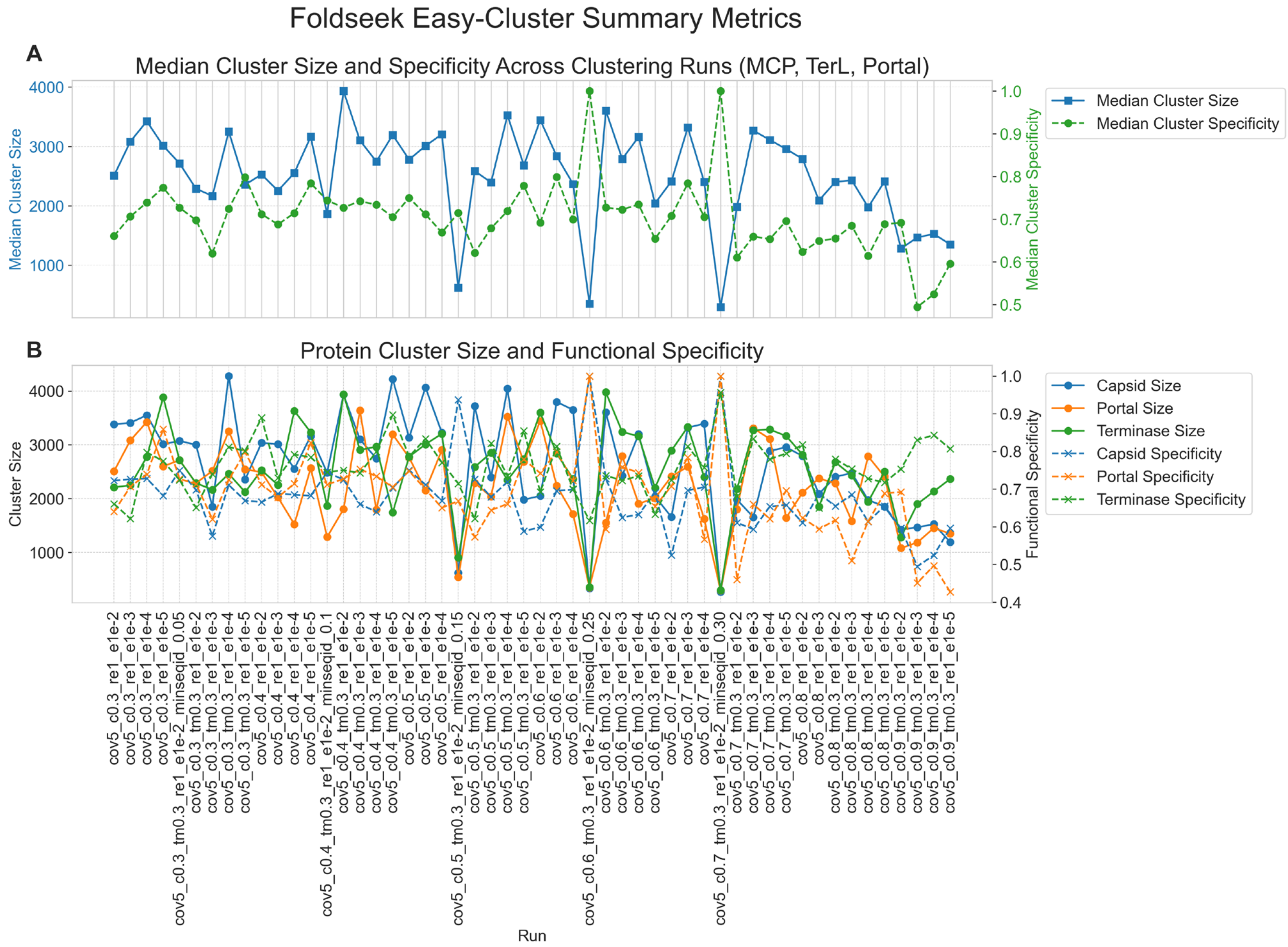


**Supplementary Figure 2. Foldseek easy-cluster summary metrics.**
**(A)** Shows the variation in median cluster size (blue, left y-axis) and median cluster specificity (green, right y-axis) for the largest clusters where the cluster representative was the major capsid protein, terminase large subunit and portal protein, across multiple Foldseek easy-cluster parameter sets. Specificity was measured as the Shannon entropy of Pfam annotations within each cluster, with higher values indicating greater functional coherence.
**(B)** Cluster sizes were tracked across three known conserved *Caudoviricetes* proteins: major capsid protein, portal protein, and terminase large subunit. As expected, high specificity is typically associated with smaller cluster sizes, reflecting a trade-off between sensitivity and specificity. To identify optimal clustering conditions, parameter sets generating a median cluster size (across MCP, portal and TerL proteins) ≥ 3,000 and a functional specificity ≥ 0.7 were selected as candidate runs for further analysis.

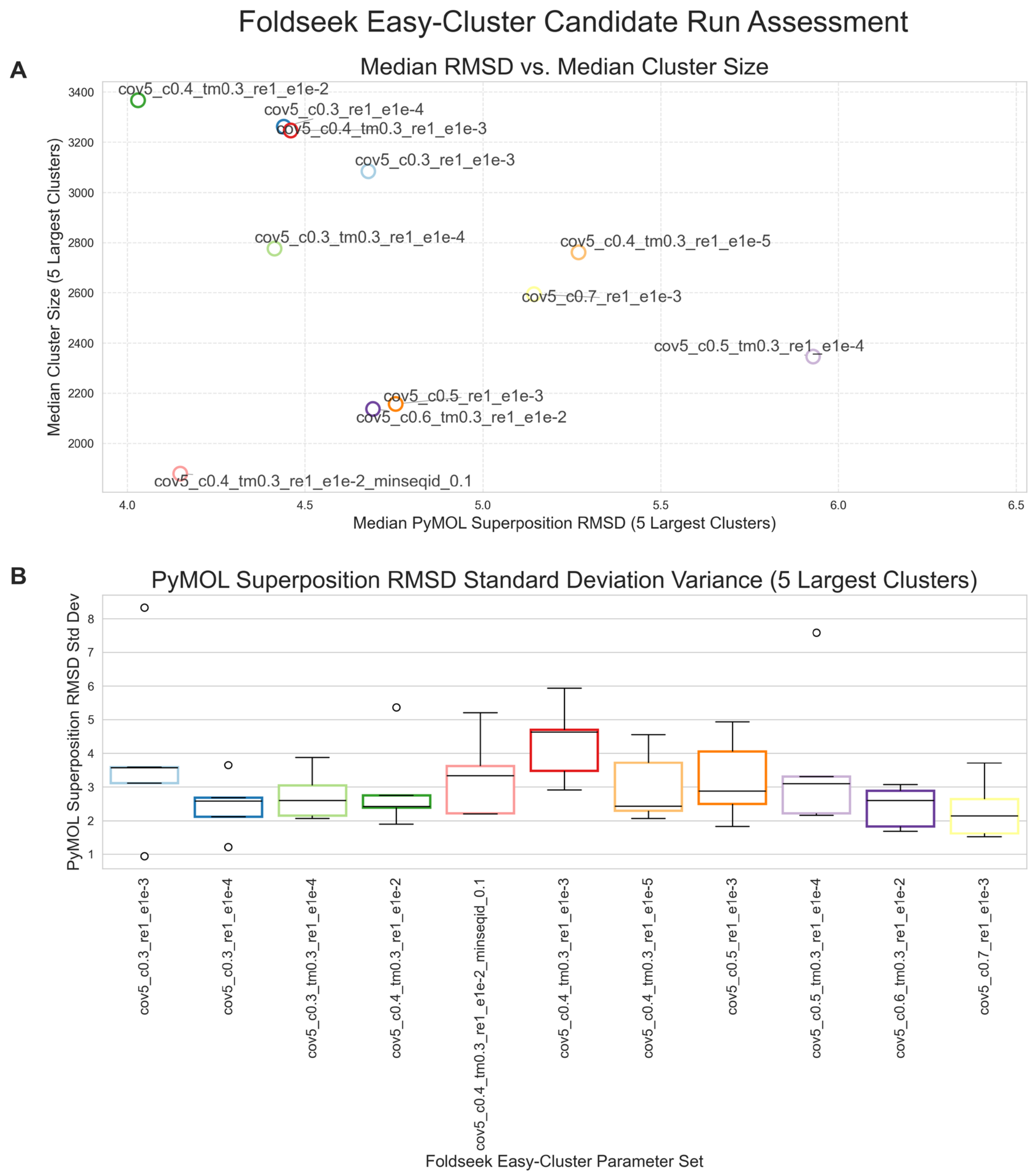


**Supplementary Figure 3. Structural coherence of Foldseek clustering parameter sets.**
**(A)** Scatterplot showing the relationship between median cluster size (y-axis) and median RMSD of PyMOL structural superpositions (x-axis) across the five largest clusters for each candidate Foldseek easy-cluster parameter set (determined by median cluster size ≥ 3,000 and a functional specificity ≥ 0.7, Supplementary Figure 2). Lower RMSD values indicate compact or high structural similarity within clusters.

**(B)** Boxplot showing the variance (standard deviation) in RMSD among members of the five largest clusters across parameter sets. Lower variance indicates greater internal structural consistency within clusters. Together, these metrics were used to evaluate the trade-off between cluster inclusiveness and structural coherence, guiding the selection of the most optimal clustering parameters: 40% coverage (mode 5), 30% TM-score threshold, cluster-reassign set to 1, and an E-value cut-off of 0.01 (1e-2).

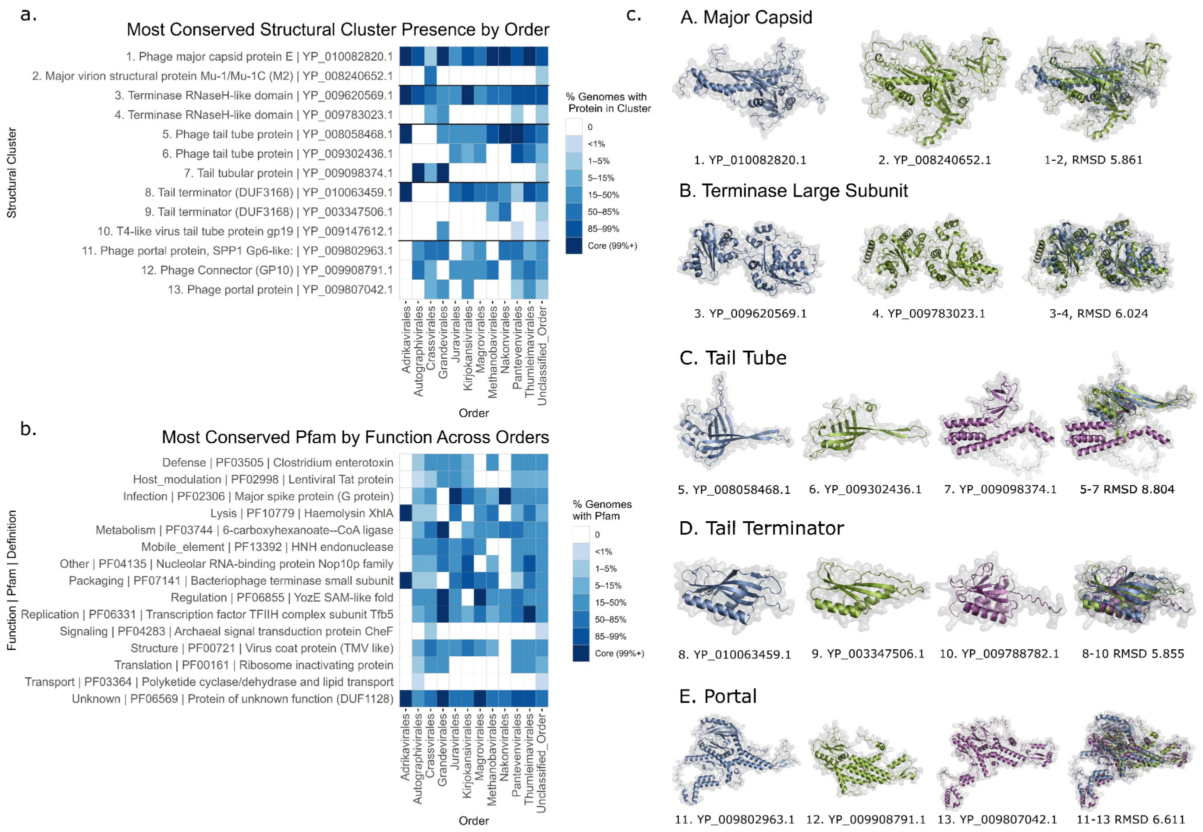


**Supplementary Figure 4. Conserved structural proteins within *Caudoviricetes* orders.**
**(a)** Heatmap showing the most conserved structural protein clusters across viral orders (ICTV VMR40) within *Caudoviricetes*, highlighting key virion components such as major capsid protein, terminase large subunit, tail tube, tail terminator, and portal proteins.
**(b)** Heatmap of the most frequently conserved Pfam domains (highest median abundance), grouped by functional category, across the same set of orders.
**(c)** Structural superpositions of protein cluster representatives belonging to the same protein family are shown in panel (a), illustrating the natural variation within each structural superfamily. Superimposition RMSD values are provided between selected pairs of cluster representatives, reflecting the degree of structural similarity among related proteins within the same functional family. When more than two proteins are superimposed, the RMSD reflects the average structural deviation between each protein homolog and the representative of the most populous cluster.

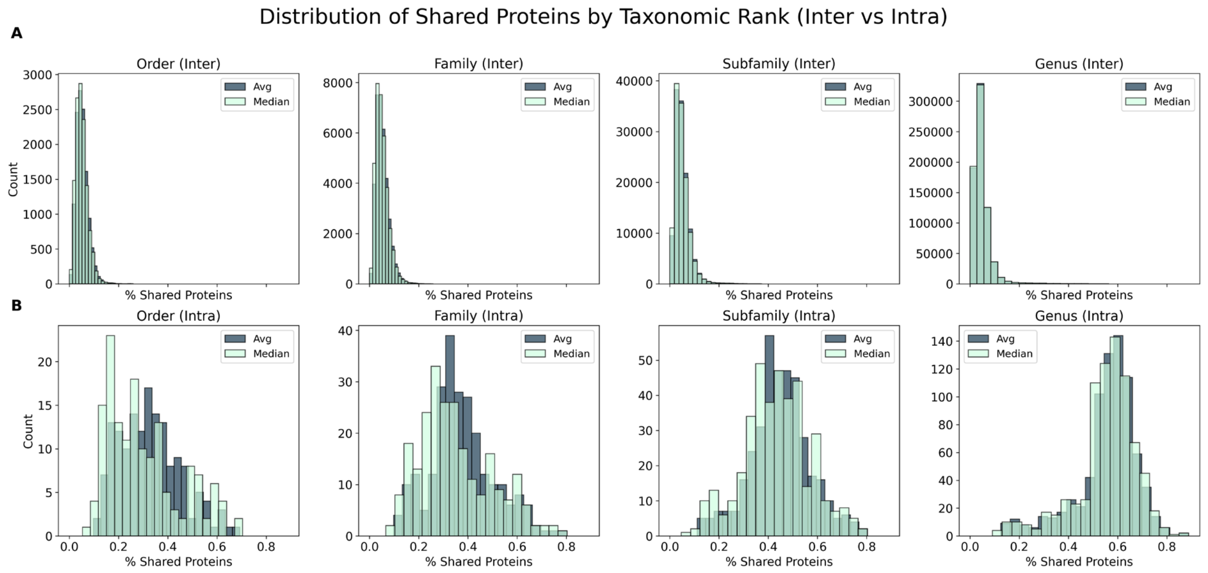


C

#### Summary of Intra- and Inter-clade Shared Protein Percentage by Taxonomic Rank

| Rank | Mean % Shared Proteins (Intra) | Median % Shared Proteins (Intra) | SD of % Shared Proteins (Intra) | Mean % Shared Proteins (Inter) | Median % Shared Proteins (Inter) | SD of % Shared Proteins (Inter) |
|---|---|---|---|---|---|---|
| Order | 0.328001 | 0.302091 | 0.121165 | 0.052271 | 0.050807 | 0.014774 |
| Family | 0.374887 | 0.350117 | 0.105864 | 0.051764 | 0.050727 | 0.012236 |
| Subfamily | 0.449797 | 0.433998 | 0.084364 | 0.049835 | 0.049204 | 0.008734 |
| Genus | 0.556628 | 0.552867 | 0.034579 | 0.048001 | 0.049204 | 0.005191 |

**Supplementary Figure 5. Distribution of shared proteins within and between taxonomic ranks.**
**(A)** Pairwise percentage of shared structural proteins between clades (inter-clade) is low across all taxonomic ranks, with narrow distributions centered around 5%.
**(B)** In contrast, intra-clade shared protein percentages are markedly higher and more variable, with distributions shifting upward at lower ranks, consistent with higher relatedness.
**(C)** Summary table showing the average, median, and standard deviation of intra- and inter-clade shared protein percentages for each rank. Intra-clade similarity increases and variance decreases toward the genus level, while inter-clade similarity remains uniformly low across ranks (~5%).

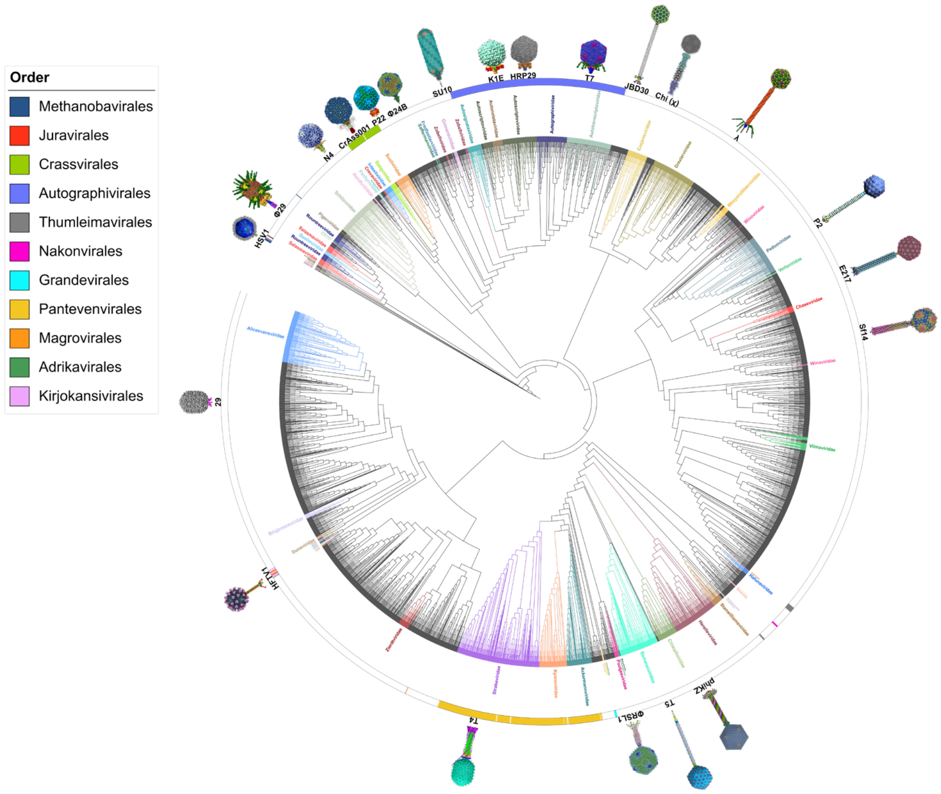


**Supplementary Figure 6. Structure-based cladogram of *Caudoviricetes* annotated with ICTV VMR40 taxonomy and cryo-EM-resolved representatives**. A phylogram constructed from pairwise Dice distances based on the presence–absence of structural protein clusters; with the topology normalized such that all branches have equal length to enhance visual clarity. Unnormalized branch lengths, reflecting pairwise Dice distances between genomes, are shown in Figure 2.

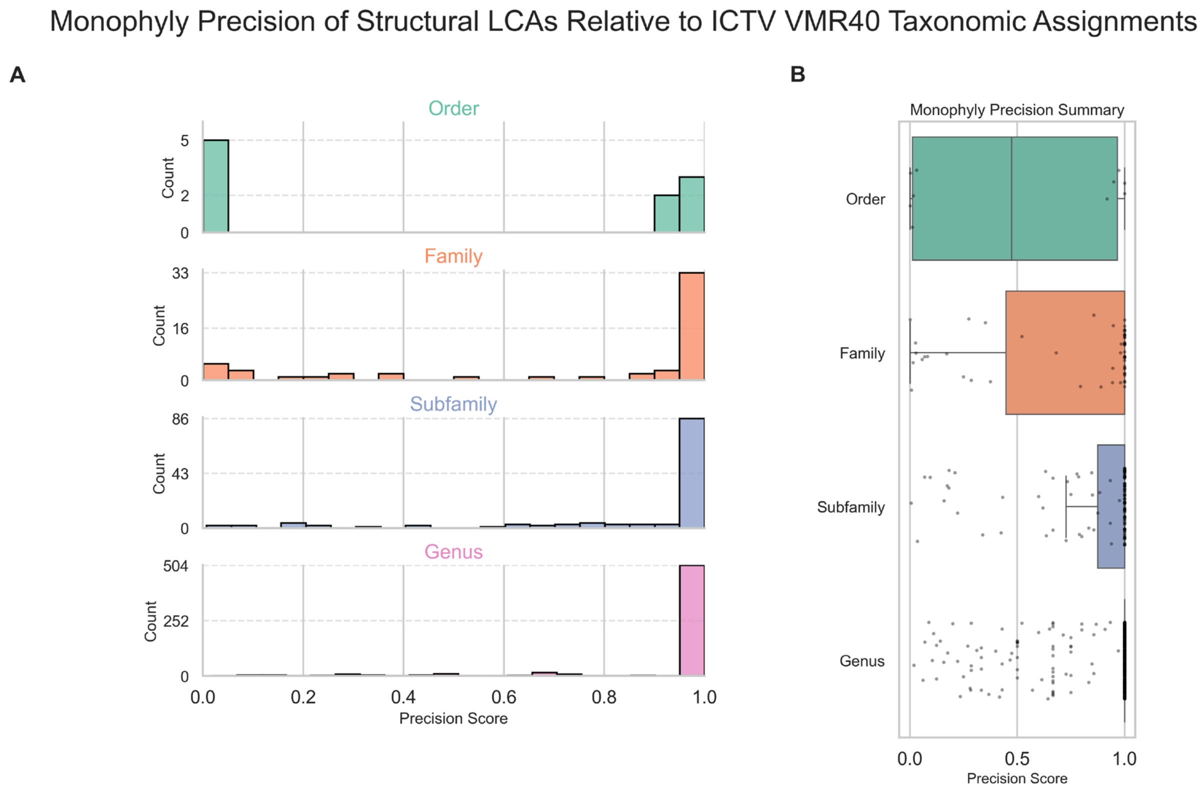


**Supplementary Figure 7. LCA-Based Monophyly Precision Across Taxonomic Ranks.**
**(A)** Histograms showing the distribution of monophyly precision scores for viral orders**,** families**,** subfamilies, and genera**.** Precision is defined as the proportion of genomes within the inferred last common ancestor (LCA) that belong to the same ICTV-assigned taxon. High precision indicates a structurally consistent and monophyletic grouping.
**(B)** Boxplots summarizing the range of precision scores across taxonomic ranks, with overlaid scatter points showing individual taxa. Overall, genus- and subfamily-level classifications show strong agreement with structure-based clustering, with the majority achieving perfect monophyly (precision = 1.0). In contrast, family- and order-level taxa exhibit greater variability, reflecting broader diversity and occasional polyphyletic placements. These scores informed subsequent RED-based reassignment of discordant clades.

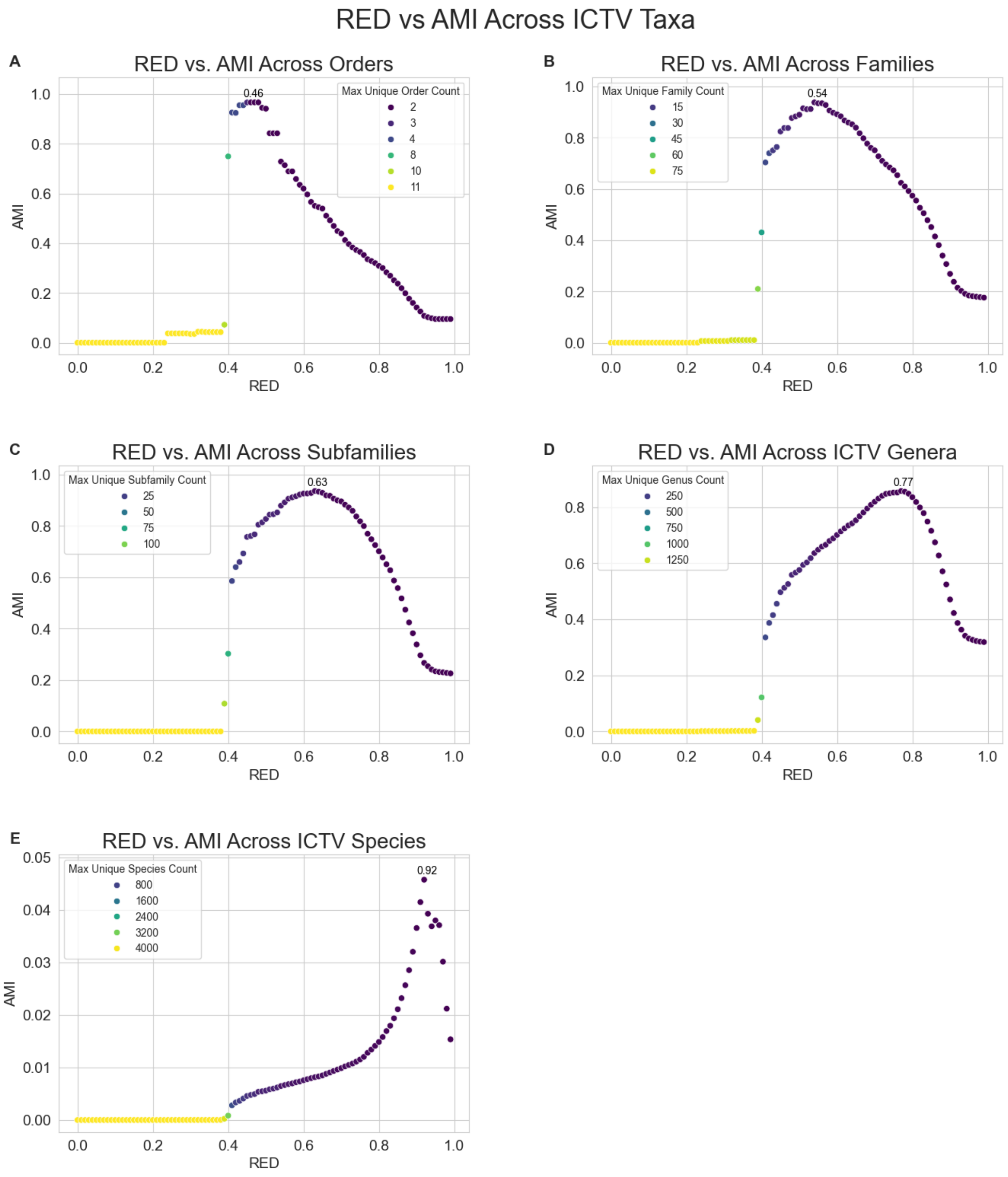


**Supplementary Figure 8. RED vs. Adjusted Mutual Information (AMI) Across ICTV Taxonomic Ranks.** Relationship between Relative Evolutionary Divergence (RED) and Adjusted Mutual Information (AMI) across ICTV-defined taxonomic ranks within the *Caudoviricetes* dataset. Each panel represents a taxonomic level—(A) Order, (B) Family, (C) Subfamily, (D) Genus, and (E) Species—and shows how well structural clustering (based on the presence–absence of structural clusters per genome) aligns with ICTV taxonomy as a function of RED score. Points are coloured by the maximum number of unique taxa assigned at each RED value for the corresponding rank.

Peak AMI values were used to define optimal RED intervals for each rank, indicating where structural coherence is highest:

- **Order:** RED = 0.46–0.54
- **Family:** RED = 0.54–0.63
- **Subfamily:** RED = 0.63–0.77
- **Genus:** RED = 0.77–0.92
- **Species:** RED = 0.92–1.00

These intervals were subsequently used as RED-informed thresholds for rank standardization in downstream taxonomic assignments.

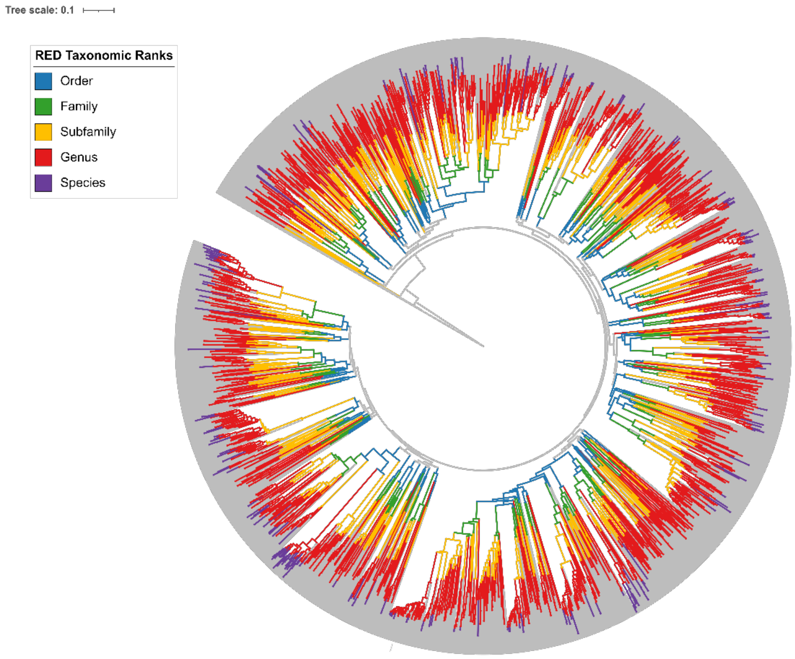


**Supplementary Figure 9. RED-Scaled Phylogeny of *Caudoviricetes* with RED vs. AMI Inferred Taxonomic Boundaries.** Circular phylogeny of 4,082 Caudoviricetes genomes, rooted on *Herpes simplex virus 1* and scaled by Relative Evolutionary Divergence (RED) using PhyloRank. The phylogram input to PhyloRank was generated from a Dice-distance structural presence–absence matrix as input (Figure 2). Taxonomic boundaries were inferred from RED intervals optimized using Adjusted Mutual Information (AMI) scores (see Supplementary Figure 6).

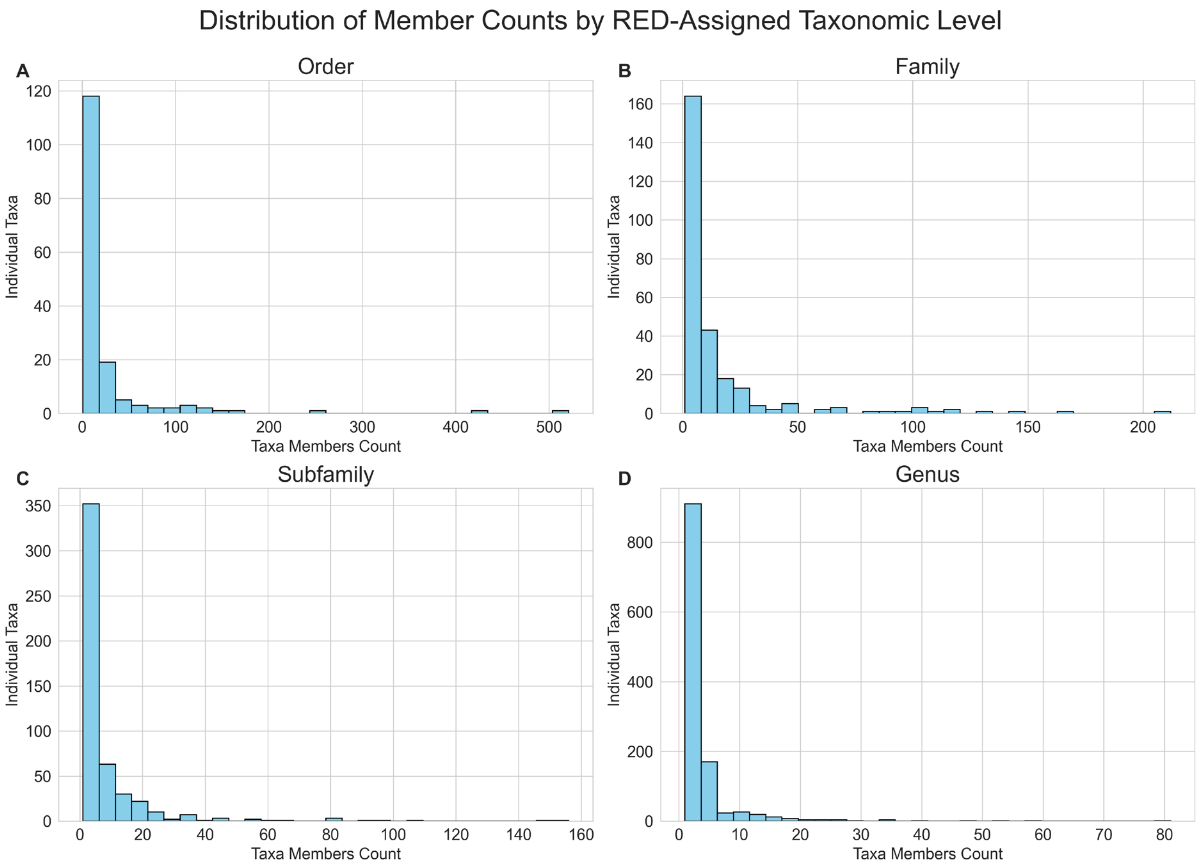


**Supplementary Figure 10. Distribution of member counts by RED-assigned taxonomic rank.** Histograms show the distribution of the number of genomes assigned to each RED-defined taxon at the order, family, subfamily, and genus levels. Most taxa across all ranks contain relatively few members, with a strong skew toward small clades and a long tail of larger taxa. At each level, the majority of taxa are composed of fewer than 10 genomes, particularly at the genus and subfamily levels, where singletons and small clusters dominate. A few large clades persist across all ranks, consistent with uneven sampling of well-studied viral groups. This distribution highlights the granularity of RED-based taxonomic assignments and the extent of diversity within *Caudoviricetes.*

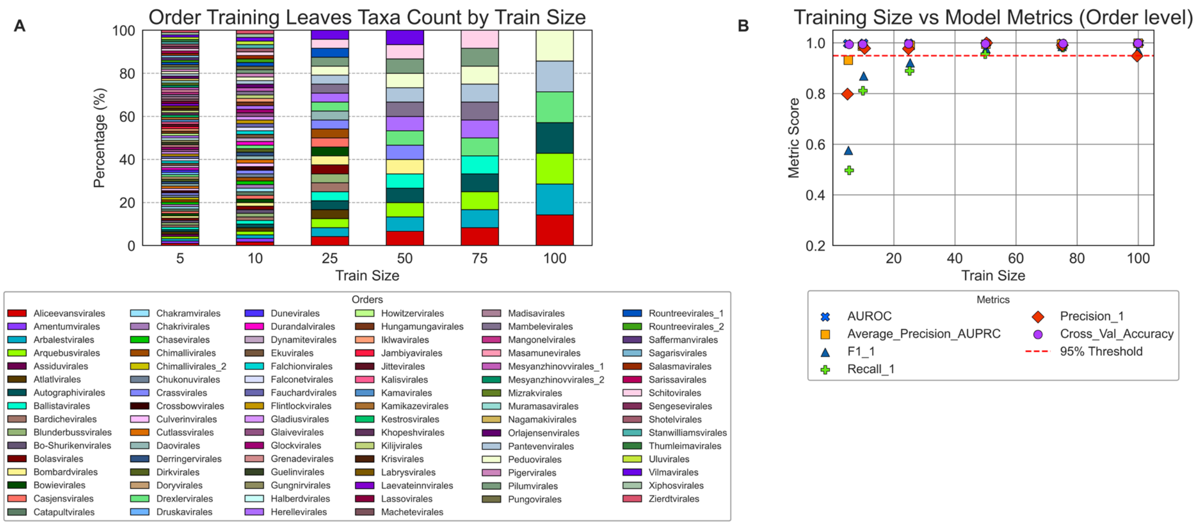


**Supplementary Figure 11. Effect of training set size on order-level taxonomic classification performance.**

**(A)** Proportion of training genomes per viral order used in binary one-vs-all models at varying training sizes (5, 10, 25, 50, 75, 100). To increase prediction difficulty and ensure fair evaluation, all taxa were downsampled to fixed sizes across orders in each iteration, standardizing the number of training taxa and preventing models from leveraging imbalanced representation. Orders with fewer than five genomes were excluded due to the requirement for 5-fold cross-validation.

**(B)** Mean performance metrics across all one-vs-all order models at each training size. Cross-validation accuracy and precision consistently remained above the 95% threshold across all sizes, and all metrics—including AUROC, AUPRC, F1, precision, and recall—exceeded 95% performance when ≥50 genomes were used for training. While the training set was downsampled to balance taxa counts, the test set was left unstratified to reflect the natural class distribution.

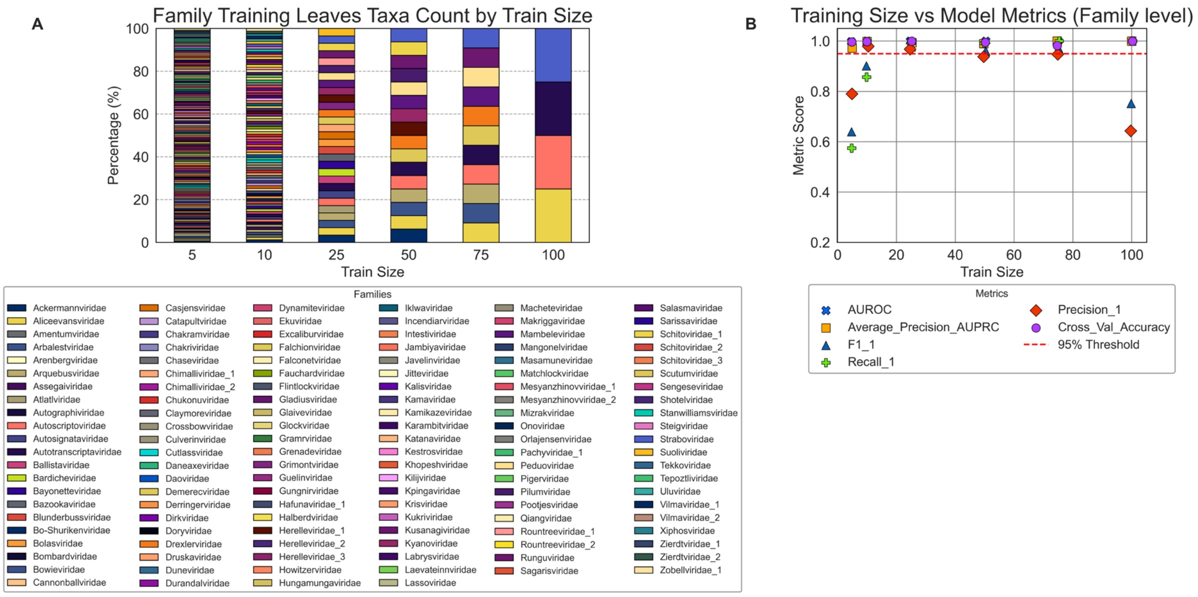


**Supplementary Figure 12. Effect of training set size on family-level taxonomic classification performance.**

**(A)** Proportional representation of training genomes per viral family in binary one-vs-all models across varying training sizes (5, 10, 25, 50, 75, 100). To standardize training conditions and increase model difficulty, taxa were randomly downsampled to equal training sizes in each round, ensuring fair comparisons across families. Families with fewer than five members were excluded to enable 5-fold cross-validation.

**(B)** Mean model performance metrics across all family-level classifiers at each training size. Most metrics, including AUROC, AUPRC, and cross-validation accuracy, remained consistently high across all train sizes. However, a decline in F1 score and precision was observed specifically at the largest training size (100), likely due to an increase in class imbalance in the test set—caused by fewer taxa being left for testing—leading to overfitting or reduced generalization capacity in these binary models when genomes seen during training were excluded.

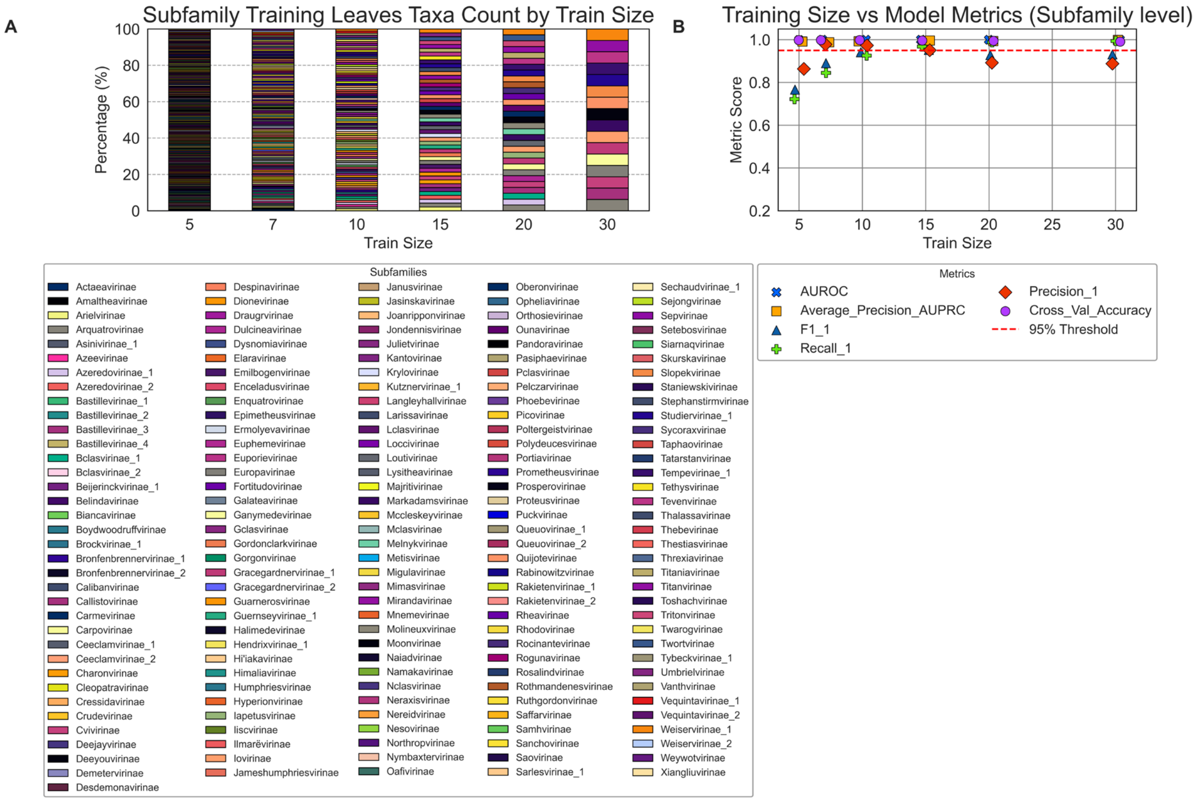


**Supplementary Figure 13. Effect of training set size on subfamily-level taxonomic classification performance.**

**(A)** Proportional representation of training genomes per viral subfamily in binary one-vs-all models across varying training sizes (5, 7, 10, 15, 20, 30). To standardize training conditions and increase model difficulty, taxa were randomly downsampled to equal training sizes in each round, ensuring fair comparisons across families. Subfamilies with fewer than five members were excluded to enable 5-fold cross-validation.

**(B)** Mean performance metrics for all subfamily-level classifiers at each training size. Performance remained consistently high for training sizes ≥10, with AUROC, AUPRC, and cross-validation accuracy all exceeding 90%. Notably, training with 15 genomes yielded the highest overall accuracy, with all metrics surpassing 95%, indicating that subfamily-level models are both robust and highly effective even at moderate training sizes.

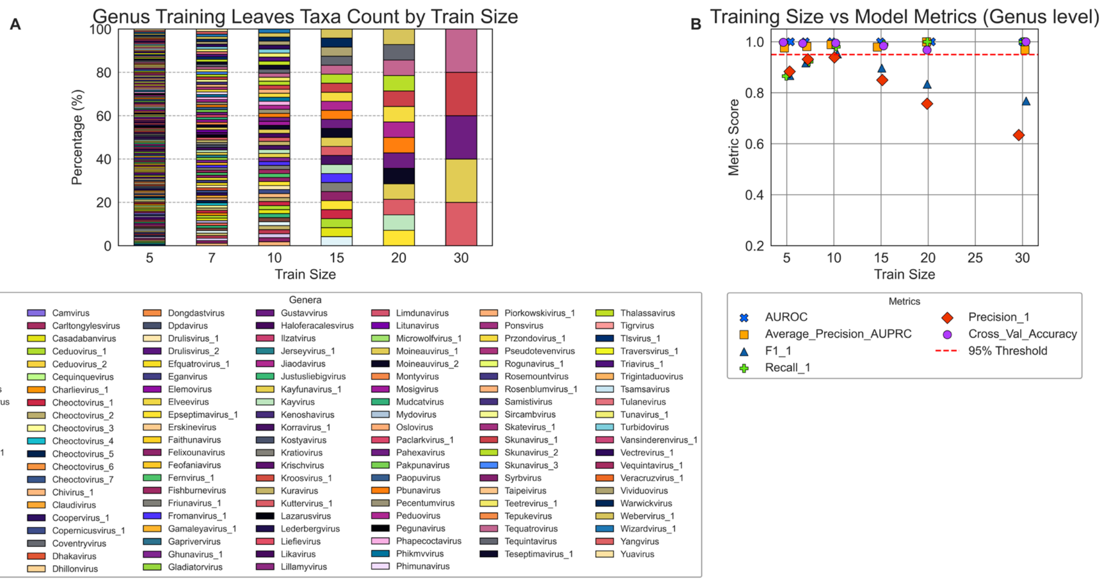


**Supplementary Figure 14. Effect of training set size on genus-level taxonomic classification performance.**

**(A)** Proportional representation of training genomes per viral genera in binary one-vs-all models across varying training sizes (5, 7, 10, 15, 20, 30). To standardize training conditions and increase model difficulty, taxa were randomly downsampled to equal training sizes in each round, ensuring fair comparisons across families. Subfamilies with fewer than five members were excluded to enable 5-fold cross-validation.

**(B)** While AUROC, AUPRC, and cross-validation accuracy remained stable, Precision_1 and F1_1 declined gradually at training sizes ≥15. This is likely due to reduced representation of the positive class in the test set which were present during training but not testing, therefore resulting in inflated false positives. The model, trained on many positives, overpredicts their presence, reducing precision and thus lowering the F1 score despite otherwise strong performance at lower train sizes e.g. 7 and 10 genomes (all metrics >=90%).

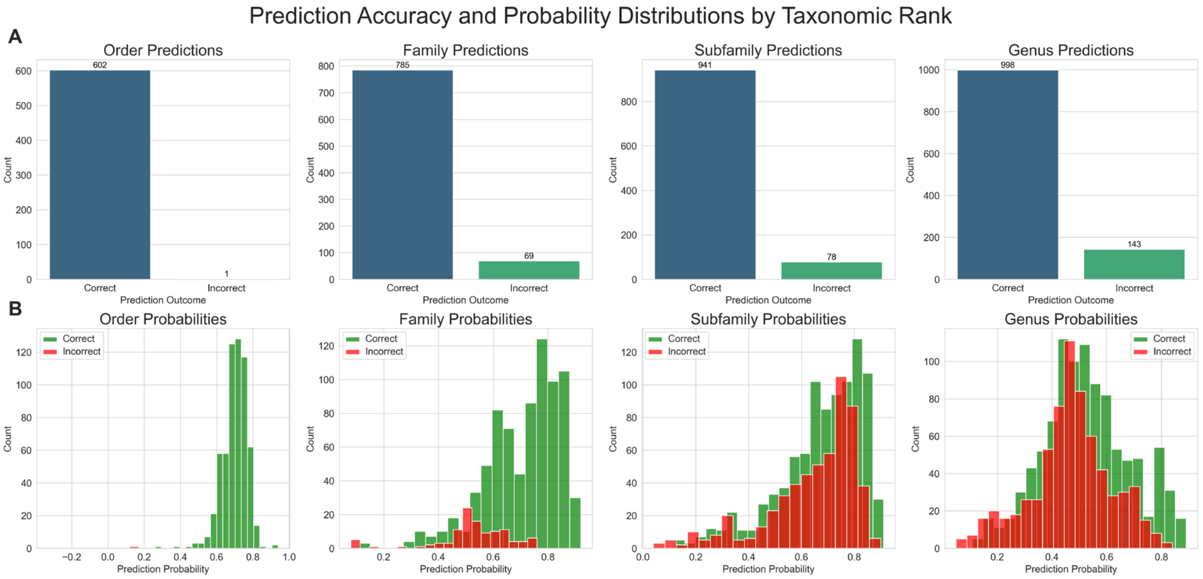


**Supplementary Figure 15. Prediction accuracy and probability distributions of the random forest classifier across four taxonomic ranks (Order, Family, Subfamily, Genus) on held-out ICTV-classified *Caudoviricetes* genomes.**

**(A)** Bar plots showing the number of correctly (blue) and incorrectly (green) classified genomes at each taxonomic rank. Classification accuracy was highest at the order level (602/603, 99.8%), followed by the family level (785/854, 91.9%), subfamily level (941/1019, 92.4%), and genus level (998/1141, 87.4%).

**(B)** Histograms of classifier probability scores for correct (green) and incorrect (red) predictions across taxonomic ranks. Correct predictions consistently exhibit higher confidence (right-skewed distributions), particularly at the order, family, and subfamily levels. In contrast, genus-level probability scores are more symmetric and broadly distributed, reflecting reduced model confidence at this rank—likely due to the limited number of training genomes per genus.

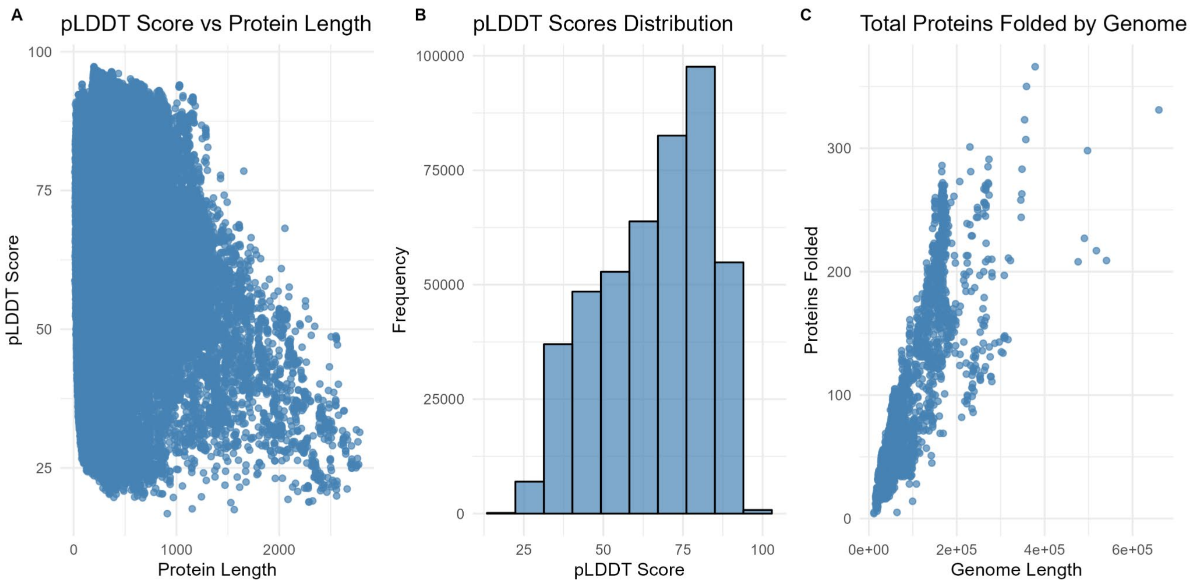


**Supplementary Figure 16. Distribution of pLDDT scores, the relationship between pLDDT scores and protein length, and the total number of folded proteins as a function of genome length.**
**(A)** Scatter plot of predicted pLDDT scores vs. protein length shows that confidence in structural predictions generally declines with increasing protein length, with shorter proteins more likely to have higher pLDDT values.
**(B)** Distribution of pLDDT scores across all predicted structures indicates a skew toward higher confidence, with most structures falling between 70 and 90.
**(C)** Total number of folded proteins per genome as a function of genome length reveals a positive correlation, with larger viral genomes encoding more proteins that could be structurally modelled.
